# Identifying Prognostic Cell State Interactions in the Tumor Microenvironment of IDH-Mutant Gliomas Using CSI-TME

**DOI:** 10.1101/2024.10.29.620901

**Authors:** Arashdeep Singh, Bharati Mehani, Vishaka Gopalan, Kenneth Aldape, Sridhar Hannenhalli

**Affiliations:** Cancer Data Science Laboratory, National Cancer Institute, National Institutes of Health Bethesda, MD, USA; Laboratory of Pathology, National Cancer Institute, National Institutes of Health Bethesda, MD, USA

## Abstract

Intercellular communication between distinct transcriptional states of various cell types in the tumor microenvironment (TME) influence the progression and clinical outcome of the tumors. Details of such clinically relevant cellular state interactions (CSIs) in IDH-mutant gliomas remain obscure. Here we developed CSI-TME, a computational pipeline that deconvolves cell type-specific gene expression from bulk transcriptomic data, identifies distinct cell states, and uncovers prognostic cell state interactions by modelling the clinical data based on the joint activity of cell state pairs. We found a significantly reproducible cell state interaction network (CSIN) that is (i) predominantly pro-tumor, (ii) likely mediated through the interaction between cell surface molecules, (iii) differentially activated in IDH-mut astrocytoma and oligodendroglioma, and (iv) significantly associated with response to immune checkpoint blockade therapy. The distinct malignant cell states involved in CSIs resembled neuronal lineages such as astrocyte-like and oligodendrocyte progenitor cells-like states and captured key interactions between glioma stem cells and immune cells. Integration of CSIN with somatic mutation data suggests the anti-tumor role of CSIs in the early stages that transitions towards pro-tumor role as glioma progresses. In summary, CSI-TME provides valuable insights into the physiology of the TME in IDH-mutant glioma and provides a framework to prioritize ligand-receptor interactions or patients stratification for therapeutic interventions.

## Introduction

Diffuse gliomas with frequent mutations in isocitrate dehydrogenase (*IDH1* or *IDH2*) enzymes (IDH-mut) have distinctive metabolic, molecular, and histological features as compared to the non-IDH mutated counterparts. IDH-mut gliomas are further classified into two categories, namely; “Astrocytoma, IDH-mutant” (IDH-A) and “oligodendroglioma, IDH-mutant and 1p/19q-codeleted” (IDH-O) (Reuss 2023; Whitfield and Huse 2022). Although these tumors respond favorably to the standard of care therapies involving maximally safe resection, radiation, or chemotherapy (Alshiekh Nasany and de la Fuente 2023), tumor recurrence is observed in almost 50% of the cases (Miller et al. 2019) and in certain cases, tumors may progress to a more aggressive secondary glioblastoma. Alternative treatment regimens such as immunotherapy have shown promising results in some clinical trials (Alshiekh Nasany and de la Fuente 2023; Gallus et al. 2023) but fail in others due to inter-tumor heterogeneity and lack of appropriate biomarkers for patient stratification (Qazi et al. 2017). Thus, there is a critical need for uncovering additional mechanisms contributing to the tumor heterogeneity and therapy resistance.

Tumor microenvironment (TME) is an ecosystem of distinct immune and non-immune cells along with the tumor cells (de Visser and Joyce 2023) and can significantly alter tumor physiology (Bremnes et al. 2011; Qu et al. 2019). For instance, despite similar expression profiles and developmental hierarchy of glial cells, IDH-A and IDH-O tumors have significantly different TME and prognosis (Venteicher et al. 2017). Further, tumor cells (Neftel et al. 2019) as well as non-tumor cells such as myeloid cells (Abdelfattah et al. 2022) and macrophages (Batchu et al. 2023) can exist in multiple transcriptional states in the TME influencing disease dynamics and therapy response. Recent studies have highlighted that the interactions between distinct cell types in the TME mediated via cell surface molecules as well as paracrine factors influence the clinical course as well therapy response in gliomas (Sharma et al. 2023). T cell-dysfunction via PD1-PDL1 interaction between T cells and tumor cells (Han, Liu, and Li 2020) and immune exclusion via TGF-beta signaling from cancer associated fibroblasts are two well-known examples of such interactions (Batlle and Massagué 2019). Additionally in glioblastoma, the secretion of interleukin-10 by *HMOX1*+ myeloid cells state can derive T-cell exhaustion (Ravi et al. 2022).

Single cell transcriptomic profiling has enabled systematic identification of cellular crosstalk by analyzing the expression of complementary ligand-receptor pairs (LR pairs) expressed by two different cell types (Armingol et al. 2021). However, there are several key limitations. Firstly, they rely on a known database of LR pairs, which are far from complete. Furthermore, cell communication might involve the coordination between distinct biological processes across cell types, either without known mediating L-R pairs, or in some cases, via other mechanisms, such as tumor-secreted metabolites (Ganjoo et al. 2023; Xia et al. 2021). Second, single cell cohorts are typically small and often lack matching clinical information of the patients that hinder their clinical assessment. Third, without appropriate controls of matched normal samples, these methods can miss the identification of interactions whose loss underlie tumor initiation.

In this work, we develop a data driven computational pipeline (CSI-TME) to infer clinically relevant cellular crosstalk between specific transcriptional states of distinct cell types in the tumor microenvironment (TME), by adapting the concept of genetic interactions (Lee et al. 2018) (Figure 1). A widely used example of genetic interactions is synthetic lethality where the joint inactivity of pairs of genes, but not individual genes, reduce cell viability (O’Neil, Bailey, and Hieter 2017). Through data driven approaches, transcriptomic and clinical datasets from large cohorts of cancer patients (such as TCGA) have been harnessed to identify synthetic lethal gene pairs (Lee et al. 2018) or more broadly gene pairs whose joint activity or inactivity significantly associated with patient survival (Magen et al. 2019). Given that tissue function is better abstracted in terms of transcriptional states on various constituent cell types, instead of individual genes, CSI-TME extends the notion of clinically relevant association between gene activity/expression to identifying clinically relevant associations between transcriptional states of different cell types in the TME. Applying CSI-TME to 425 IDH-mut glioma RNA-seq samples from TCGA, we identify a robust and reproducible network of transcriptional state interactions across various cell types. A majority (70%) of these interactions were pro-tumor, with a minority (30%) having anti-tumor associations. CSI-TME identified interactions that involved known states of malignant cells such as astrocyte-like, oligodendrocyte progenitor-like, and cycling stem cells, and provided a framework to prioritize ligand-receptor interactions with their predicted clinical outcomes. Integrating the identified CSIs with paired somatic mutation data, we observed that despite their overall under-representation in the CSIN, anti-tumor interactions were significantly over-represented in the samples bearing somatic mutations in IDH-mut glioma patients raising the possibility that TME responds to somatic mutations to restore tissue homeostasis. The differential enrichment of anti-tumor interactions among the mutated samples in early stages relative to later stages of tumorigenesis further supported this hypothesis. Overall, implementing a novel computational strategy, CSI-TME provides new insights into the cellular interactions in the IDH-mut glioma TME with a potential application in patients stratification for therapeutic interventions. The open-source software for CSI-TME is available at https://github.com/hannenhalli-lab/CSI-TME.

**Figure 1.**
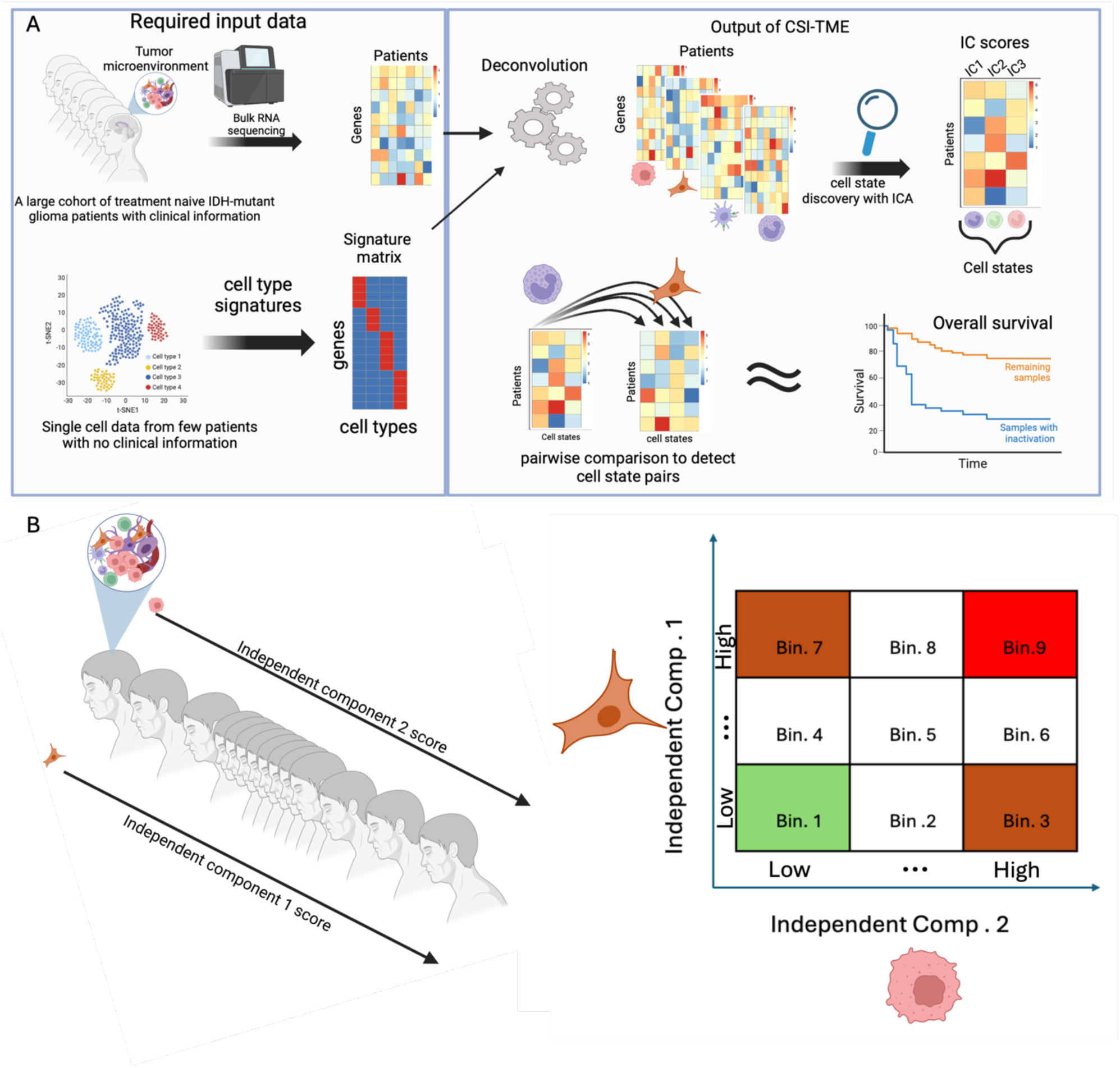
A schematic of the key steps involved in CSI-TME. **A** Given (i) bulk transcriptomic data from a large cohort of patients along with survival data and (ii) cell type specific gene markers inferred from single cell data, CSI-TME infers cell state pairs across different cell types that are significantly prognostic based on their joint activity status. Following deconvolution, each cell-type specific gene expression matrix is subjected to ICA; the resulting independent components are deemed to quantify the abundance of distinct cell states across patients (cell state matrix). Finally, we perform a Cox-regression to model the overall survival of patients based on the joint activity IC-pairs taken from two different cell types. **B** For any two pairs of ICs, CSI-TME considers their joint activity in following three categories: both ICs are low (Bin1), both ICs are high (Bin9) and either of the IC is low and other IC is High (Bin3 and Bin7)

## Results

### An overview of CSI-TME

A schematic of the key steps involved in CSI-TME is illustrated in Figure 1 and the details are provided in Methods. Briefly, given a large cohort of bulk tumor transcriptome (RNA-seq) data and cell type specific marker genes for key cell types, CSI-TME first deconvolves the bulk expression data into their constitutive cell type-specific expression profiles using CODEFACS (Wang et al. 2022).

Next, for each cell type, CSI-TME infers their distinct cell states or gene expression programs based on independent component analysis (ICA), to obtain an IC-by-sample matrix for that cell type where each IC represent a distinct transcriptional state. Finally, CSI-TME screens for the pairwise combinations of transcriptional states (ICs) between two distinct cell types whose simultaneous activity or inactivity (for instance, senescent T cells and stem-like malignant cells) significantly associates with patient survival using Cox proportional hazard test. The output of CSI-TME is a list of tuples (C1, S1, C2, S2, B, D), where co-occurrence of transcriptional state S1 of cell type C1 and transcriptional state S2 of cell type C2 in a specific co-activity bin B is associated with survival, where D=+1 indicates positive log hazard ratio and D=-1 indicates negative log hazard ratio. For a given cell type, samples are partitioned into three equal-sized bins -- Low, Med, and High – based on the score of the IC. We consider three co-activity bins (Figure 1B) corresponding to co-inactivity of S1 and S2 (bin 1), co-activity of S1 and S2 (bin 9), and high activity of S1 and low activity of S2 (bin 3) with the converse symmetric case being included (i.e. bin 7, Figure 1B).

### CSI-TME identifies distinct transcriptional states and their pro- and anti-tumor interactions in the TME of IDH mutant glioma

Integrating a publicly available scRNA-seq data with a cohort of our in-house scRNA-seq data from glioma patients (Methods), with a total of 60751 *cells*, consistent with previous studies, we identified seven distinct cell types based on the expression of established cell type-specific marker genes (Figure 2A). Using these 7 cell type-specific gene signatures, we used CODEFACS to deconvolve two large cohorts of bulk gene expression data from TCGA (discovery cohort) and CGGA (validation cohort) IDH-mut gliomas, yielding seven cell type-specific expression profiles for each patient across these two cohorts (Methods), covering, on average across the cell types, 9954 genes in TCGA and 11631 genes in CGGA.

**Figure 2.**
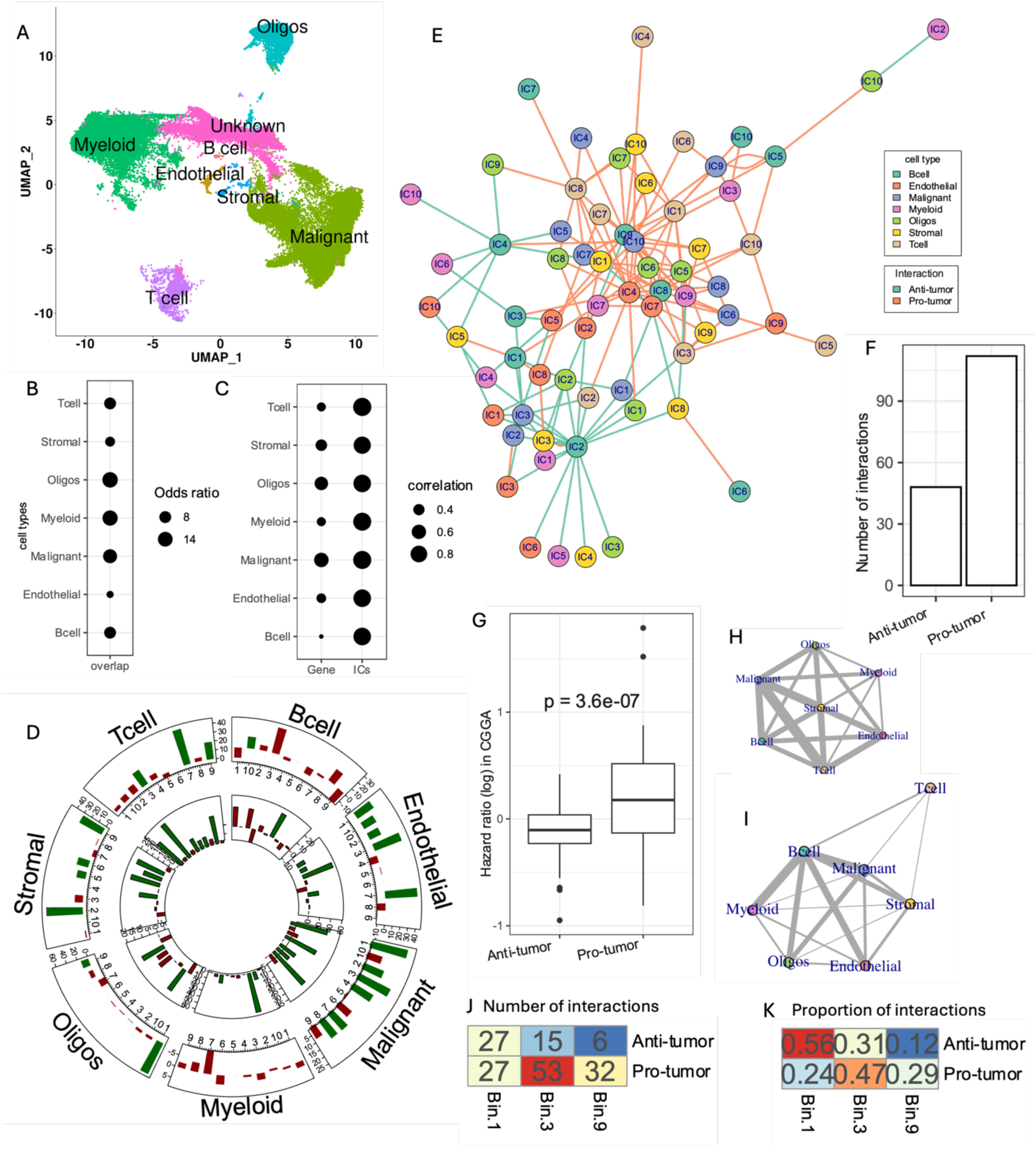
Transcriptional states and their interactions in IDH mutant gliomas. **A.** UMAP plot of the scRNA-seq data from IDH-mutant patients. Cells are colored and the cell type labels shown for each cluster. **B.** A dot plot showing the odds ratio of overlap among the top genes expressed by each cell type between TCGA and CGGA in the deconvolved data; all overlaps were significant (FDR < 0.20) based on Fishers’ test. **C.** Dot plot showing the cross-gene correlations (left column) of log hazard ratio between TCGA and CGGA. Similar correlation between independent components is shown in the right-side column. **D.** Bar plots in circular layout. Each sector corresponds to a distinct cell type as labeled, and each bar represents a distinct IC (x-axis) for that cell type. All inter-cell distances (Euclidean distance in PC space) were computed and partitioned into intra-IC (both cells have high IC score) and inter-IC (exactly one cell has high IC score). The height of bars denotes the percent difference between the inter-IC and the intra-IC distances such that a positive (respectively, negative) value indicates higher (respectively, lower) similarity between intra-IC cell pairs. A bar is shown in green if the inter-IC distances are significantly greater than intra-IC distances (difference > 10% and FDR < 0.20), implying the tendency of that ICs to cluster in PC space. The outer track is for positive signature genes and the inner track is for negative signature genes of the ICs. **E.** CSIN involves distinct cell states and gene expression programs derived from 7 distinct cell types. Node colors represent the cell type and edge color represent the prognostic effect of each interaction. **F.** A bar plot showing the number of pro-and anti-tumor interactions in the CSIN. **G.** Boxplots showing the hazard ratio of pro- and anti-tumor interactions in CGGA. **H-I.** CSIN summarized at the level of cell types for pro-tumor (H) and anti-tumor (I) interactions. The width of the edges is proportional to the number of interactions between each cell type pair. **J-K.** Distribution of the number of interactions across different activity bins (J) and proportion of interactions leading to better (anti-tumor) or worse (pro-tumor) prognosis in each bin (K). These plots show that interactions leading to better prognosis are enriched in Bin 1.

Before proceeding, we evaluated that deconvolution captured the cell type specific expression signals in each tumor (Figure 2B, C and Supplementary Figure S1A, B). Firstly, the top 5% highly expressed genes in each cell type in TCGA significantly overlapped with the top 5% genes expressed in the same cell type of CGGA samples (Figure 2B); in contrast, the overlap among the highly expressed genes across distinct cell types in TCGA and CGGA was significantly lower (Supplementary Figure S1B). Functional enrichment analysis of the top 5% genes expressed by each cell type revealed several cell type specific functions (Supplementary Figure S1A, Methods). For instance, immune system terms related to myeloid, and leukocytes were enriched in the myeloid cells. Functions related to the central nervous system such as axon ensheathment, neurogenesis, neuronal differential, and glial cell fate commitment were enriched among the malignant cells and OPCs. To further assess the robustness of deconvolution, we performed a Cox regression for each gene using the deconvolved gene expression data in each cell type in TCGA and CCGA (Methods) and computed the across-gene correlations of log hazard ratio (lHR) between the two cohorts. For the same cell type, the cross-gene correlation of lHR between two cohorts ranged from 0.22 to 0.6 (Figure 2C, left column). Furthermore, the cross-gene correlations of lHR were substantially higher when comparing the same cell types between TCGA and CGGA, in contrast to comparing two different cell types (Supplementary Figure S1C). Together these results support the robustness of the deconvolved data.

After deconvolution, we identified distinct transcriptional states for each of the seven cell types by using independent component analysis (ICA) to derive latent factor projections or independent components (IC) of the deconvolved TCGA data (Methods). In each cell type, we derived 10 ICs, where each IC represents a distinct transcriptional cell state or gene expression program (Methods). For validation, we projected the deconvolved data from CGGA cohort onto the latent factor space derived from TCGA cohort (Methods). Interestingly, many of the ICs were significantly prognostic in TCGA and had a significantly more consistent association with patient survival in the validation cohort (CGGA) as compared to the individual genes (Figure 2C and Supplementary Figure S1D). These results emphasize the value added by the latent factor projection of the deconvolved data in capturing the cell type specific signal.

The ICs derived from each cell type might represent distinct cell states or their gene expression programs that might be shared across cell states. We assessed the extent to which the ICs represent specific transcriptional states of the corresponding cell type in a scRNA-seq dataset of IDH-mut glioma (Methods). For this analysis, we scored the single cells in scRNA-seq data for expression of signature genes representing each IC and labelled cells that do or do not express IC signature genes (Methods). We observed that in several cases, cells expressing IC signature genes tend to be significantly closer to each other than the cells that do not express the IC signature genes in principal component space (Figure 2D). UMAP projections of some of the ICs that show significant clustering in principal component space are provided in supplementary Figure S1E-G). A total of 54 out of 70 ICs showed significant tendency for clustering across all cell types (Figure 2D). These ICs thus might represent distinct cell states of the respective cell type. The remaining ICs that do not exhibit clustering in scRNA-seq might represent other cellular programs shared across cell states. To further characterize the biological functions of ICs, we performed functional enrichment analysis (Methods) and observed that several housekeeping as well as cell type specific functions were enriched among the signature genes of ICs (Supplementary Table S2). Together, these results suggest that ICA of the deconvolved cell types captures distinct cell states, or their gene expression programs in the TME. The notion that ICs might represent cell states is further strengthened by the observation that signature genes of several ICs had a significant linear clustering on the chromosomes (Supplementary Figure S1J-K), consistent with the idea of coordinated epigenetic regulation underlying cell state switches (Madrigal et al. 2023).

Finally, we aim to identify interactions between transcriptional states (represented by the ICs in each cell type) across cell types, associated with patient survival in TCGA. Using Cox regression, we performed a computational screen for all pairs of ICs from two distinct cell types whose joint activity levels (either Low or High) were significantly associated with the survival of IDH-mutant glioma patients (Methods). For instance, we tested for assertions such as “Joint occurrence of low activity of IC1 in malignant cells and high activity of IC3 of T cells is associated with worse prognosis” or vice versa (Figure 1B). At the FDR threshold of 20% and 70% internal cross-validation accuracy (Methods), we detected 160 significant interactions, a majority (70%) of which were associated with worse prognosis (Figure 2E, F). Together, these 160 interactions form a network of CSIs that we refer to as CSIN. We term the interactions leading to worse or better prognosis as pro-tumor or anti-tumor cell state interactions, respectively. To ascertain if the detected CSIs are reproducible, we deconvolved an additional cohort of 325 IDH-mutant glioma patients from CGGA and scored each sample using the rotation matrix of the ICs derived from TCGA (Methods). We then performed an analogous detection of CSIs in the CGGA cohort. Encouragingly, a majority of the interactions that lead to better/worse prognosis (i.e. anti-tumor and pro-tumor) in TCGA tended to have a negative/positive log-hazard ratio in CGGA as well (Figure 2G). Additionally, as a negative control, we generated sets of interactions by randomizing the clinical data of patients in TCGA ten times (Methods) and observed that random interaction sets did not significantly overlap across the cohorts (Supplementary Figure S1H). These results support the robustness as well as potential biological significance of the detected interactions.

**Figure S1.**
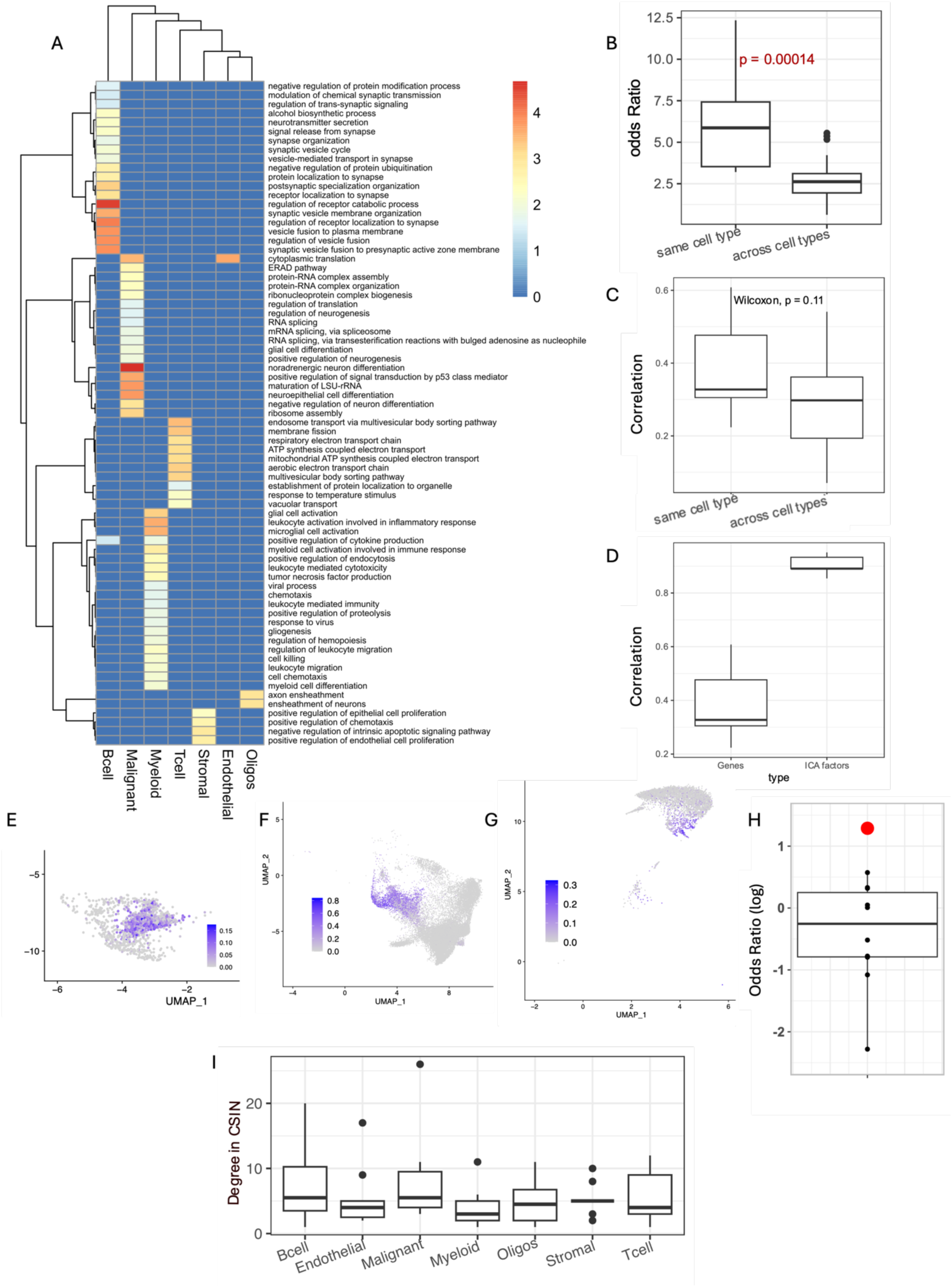
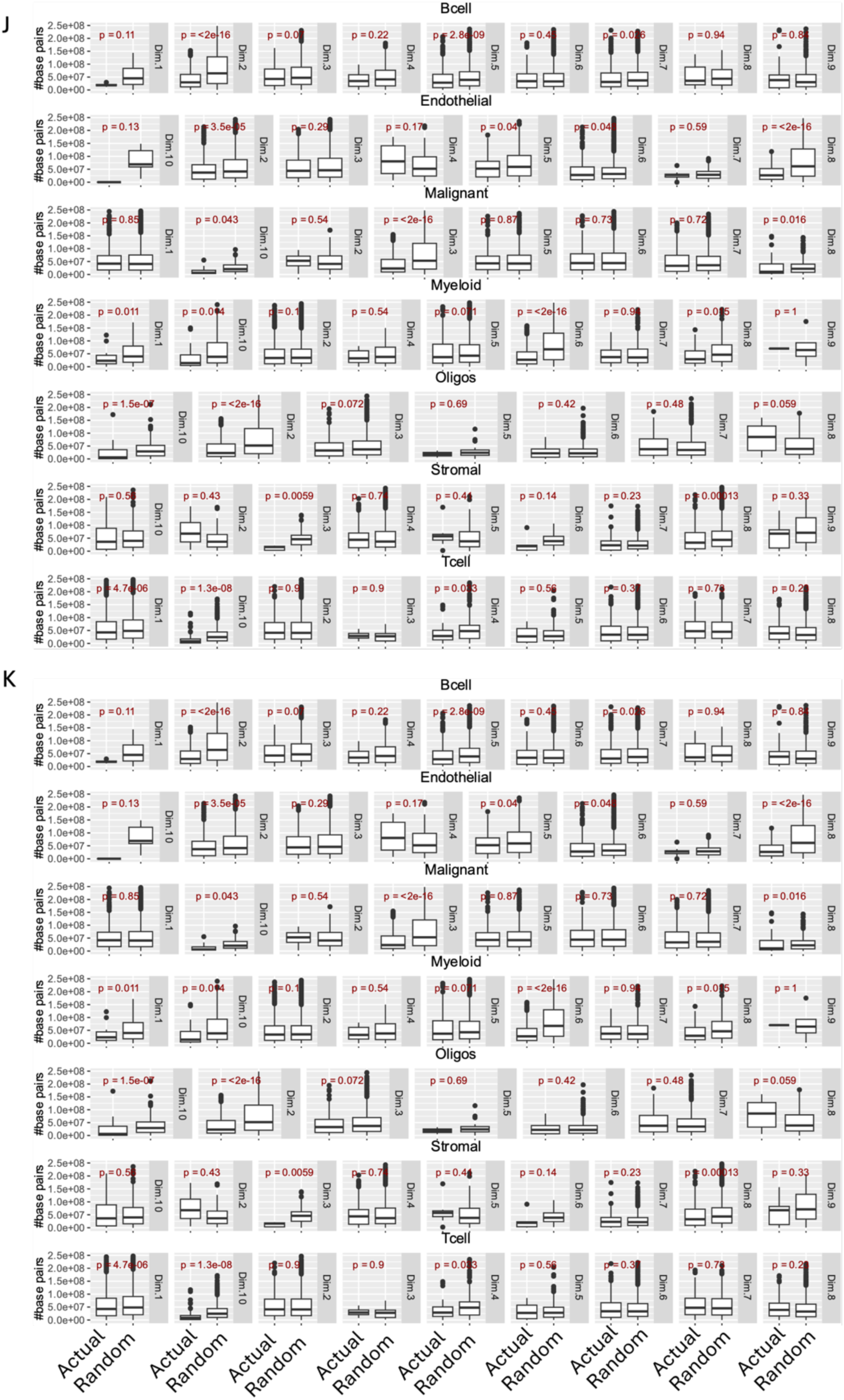
**A.** Heatmap showing the enrichment of distinct functions among the top genes expressed by each cell type in the deconvolved data. **B.** Boxplot showing the odds ratio of enrichment among the top genes expressed between the same cell type and across different cell types in the deconvolved data in TCGA and CGGA **C.** A box plot showing the cross-gene correlations (left column) of log hazard ratio between TCGA and CGGA for cell same type and across cell type comparisons. **D.** Boxplot showing the distribution of correlation of hazard ratios between same cell types computed using genes or using ICs. **E-G.** UMAP plots for T cells (E), malignant cells (F) and Oligodendrocytes (G) colored by the signature scores derived from the signature genes of IC1 (T cells), IC5 (Malignant cells), and IC7 (Oligodendrocytes), respectively. **H.** Boxplot showing the odds ratio (log scale) of enrichment for significant interactions (nominal p-value < 0.05) between TCGA and CGGA (red point) using the randomized clinical data. The odds ratio between actual interactions at a similar p-value threshold is shown by the red point. **I.** Boxplot showing the distribution of the number of interactions observed for the ICs of each cell type. **J.** Boxplots showing the linear clustering of the positive signature genes of various ICs. On Y axis is the distribution of pairwise linear chromosomal distances (in base-pairs) among the positive signature genes of various ICs (i.e. Dim.1, Dim.2 etc.) that are present on same chromosome. For each IC, the actual distribution of linear-chromosomal distances between the signature genes is compared against the distribution of genomic distances between 10 randomly drawn gene sets. These randomly drawn gene sets had a similar chromosomal distribution to the signature genes of the corresponding ICs. P-values from Wilcoxon’s tests are shown. **K.** Same as J but for negative signature genes.

In the CSIN, each cell type had a variable number of interactions (Supplementary Figure S1I) with malignant cell and B cell states involved in the greatest number of interactions. A majority of the detected interactions involving T cell states (ICs) associated with worse survival (i.e., pro-tumor) and largely involved interactions with malignant cells (Figure 2H). This is consistent with the idea of tumor cell-driven T-cell exhaustion as well as emerging pro-tumor roles of certain T-cell states (Reis et al. 2022). We probed the distribution of pro-tumor and anti-tumor interactions across their co-activity bins and found that despite a three-fold greater number of pro-tumor interactions in the CSIN, anti-tumor interactions were significantly over-represented in Bin 1 (Figure 2J-K).

### CSI-TME identifies glioma stem cells resembling neuroepithelial lineages of the developing human brain and their associated interactions in TME

Tumor cells exist in multiple transcriptional states that recur across multiple cancer types (Barkley et al. 2022). We assessed the extent to which the ICs derived from IDH-mut malignant cells capture these conserved cell states. Toward this, we assessed the significance of overlap between the signature genes of the recurring cancer cell states obtained from Barkley et al. and the signature genes of the ICs (positive as well as negative) from malignant cells (Methods). We observed that the negative genes of the three of the malignant cell ICs significantly resembled oligodendrocyte progenitor (IC5), astrocytic (IC6), and cycling cells (IC7) (Figure 3A). GO functional enrichment analysis further supported the proliferative nature of IC7 (Supplementary Figure S2A). Recent work has shown that malignant cells in IDH-mutant glioma exist in three distinct states including a highly proliferative population of glioma stem cells (GSCs) resembling the neuronal precursor cells (NPCs) which differentiate into astrocytic and glia-like oligodendrocyte lineages, thus recapitulating the developmental hierarchies (Tirosh et al. 2016). Several reports have indicated that mutation in the IDH1 gene can block differentiation and induce stemness (Haddock et al. 2022; Y. Zhang et al. 2022). Therefore, we assessed whether IC7, which captures the proliferation state of the cell (Figure 3A), represents a population of GSCs. Toward this, we first defined a set of 1143 putative stemness signature genes (PSS1) in IDH-mut glioma as the genes that were significantly positively correlated with the expression level of *IDH1* across samples in the TCGA cohort of IDH-mut glioma (Methods). We observed that PSS1 had significantly negative gene weights in IC7 (Figure 3B). PSS1 also exhibited a significant overlap with negative signature genes of IC7 (Figure 3C). To assess the robustness of this observation, we obtained an additional putative stemness signature (PSS2) derived from single cell investigations of IDH-mut gliomas from a previous study (Venteicher et al. 2017) and observed that PSS2 also had significantly negative gene weights in IC7 as well as showed a significant overlap with the negative signatures genes of IC7 (Figure 3B, C), suggesting that IC7 captures the transcriptional program in the glioma stem cells axis (Figure 3B).

**Figure 3.**
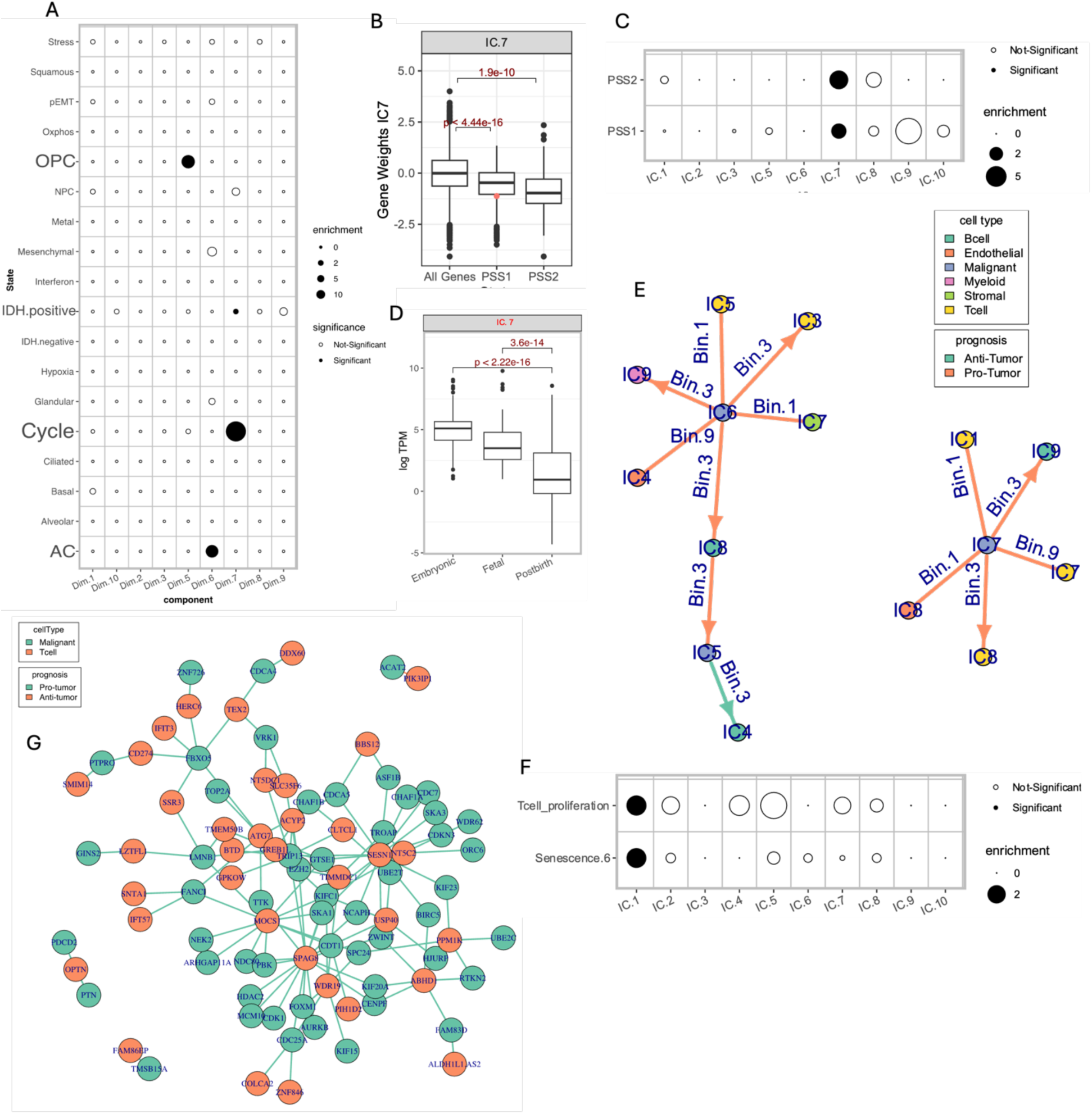
Identification of glioma stem cells and their interactions with other cell types in TME. **a.** A dot plot showing the odds ratio for the overlap between the negative signature genes of various Malignant cell IC and recurrent cancer cell states. Size of dots is proportional to the odds ratio derived from Fisher’s test and solid points indicate significance (FDR < 0.20). **b.** A box plot showing the contribution of PSS signature genes to IC7 of malignant cells. Red dot is the contribution of the *CD44* gene which is not included in the PSS. **c.** A dot plot showing the enrichment of two putative stem cell signatures (PSS1 defined in this study based on correlation with IDH1 expression, and PSS2 sourced from Venteicher et al.) among the malignant cell ICs. The shape and size of the points follows the similar convention as in panel a. **d.** A box plot showing the distribution of log normalized TPM values for the signature genes of IC.7 during different developmental stages of the human brain. **e.** A network plot of CSIs involving IC5, IC6, and IC7 of the malignant cells. **f.** A dot plot showing the odds ratio for the overlap between negative signature genes of T cells ICs against the markers for T cell proliferation (GO:0042098) and senescence (from Wechter et al.). **g.** A network plot showing the interactions between negative signature genes of IC7 of Malignant cells and IC7 of T cells.

Among PSS2, overlapping genes include a bona fide neurodevelopmental transcription factor *SOX11* (Venteicher et al. 2017). Therefore we further probed the expression pattern of the signature genes for IC7 from malignant cells using a publicly available dataset of developing human brain (Cardoso-Moreira et al. 2019) (Methods). Interestingly, the negative signature genes of IC7 had higher expression during the embryonic phases of brain development and significantly decreased in fetal brain and mature adults (Figure 3D) and no other IC showed this trend (Supplementary Figure S2B). Additionally, *CD44* which is known to be highly expressed by GSCs and has a hypoxic condition-specific effect on invasion and proliferation (Inoue et al. 2023) had a negative loading value in IC7 (Figure 3B). Together, these observations establish that the negative direction of IC7 represents a population of highly proliferative GSCs resembling developing neuronal cells.

Recent studies have shown that crosstalk between GSCs and the immune system contributes to therapy resistance (Eckerdt and Platanias 2023). Therefore, focusing on the interaction involving GSCs and the components of the immune system (Figure 3E), we observed that IC7 of malignant cells engaged in pro-tumor interactions with IC9 of B cells and IC1,7,8 of T cells spanning all three activity bins. To aid interpretation, we present a schematic table in Supplementary Figure S2C that outlines the status of negative and positive signature genes for the two interacting ICs across various interaction bins. An interaction of malignant cell IC7 (related to GSC) involving IC1 of T cells occurred in Bin 1 associated with worse survival (pro-tumor). We explored the possible functions associated with IC1 of T cells and found that negative signature genes in the IC1 of T cells showed a significant enrichment of functions related to cell cycle progression such as G1/S transition and chromosomal segregation (Supplementary Figure S2D). Next, focusing on GO term related to the proliferation of T cells, we observed that negative signature genes of T cell IC1 significantly overlapped with the genes annotated to be involved in T cell proliferation (GO:0042098) (Figure 3F), further confirming that negative loadings in IC1 of T cells represent proliferating T cells. Since the interaction in bin 1 implies the upregulation of negative signature genes of both the IC partners (Supplementary Figure S2C), it is intriguing that malignant cell stemness, together with T cell proliferative state, is associated with worse survival. Proliferation of T cells in TME can induce their senescence due to telomere shortening (Woroniecka et al. 2018). However, GSCs, which are also proliferating, have the ability to restore telomeres (Woroniecka et al. 2018). Therefore, we assessed if the negative signature genes of T cell IC1 also resembled the markers of senescence. Although systematic investigations of T cell senescence are lacking, we observed that certain lymphocyte specific markers of senescence curated from literature had negative gene-weights in IC1 (Supplementary Figure S2E). Additionally, we obtained a gene signature that is upregulated in the senescent population of cells from a systematic scRNA-seq investigation focused on fibroblasts (Wechter et al. 2023). Interestingly, we observed that negative signature genes of T cells IC1 significantly overlapped with the markers of senescent fibroblasts (Figure 3F), possibly indicating that IC1 captures an intermediate state between proliferating and replication-induced senescent cells. Hence, our analysis suggests that the synergistic effect of GSCs coupled with T cell senescence might result in the worsened patient survival.

**Figure S2.**
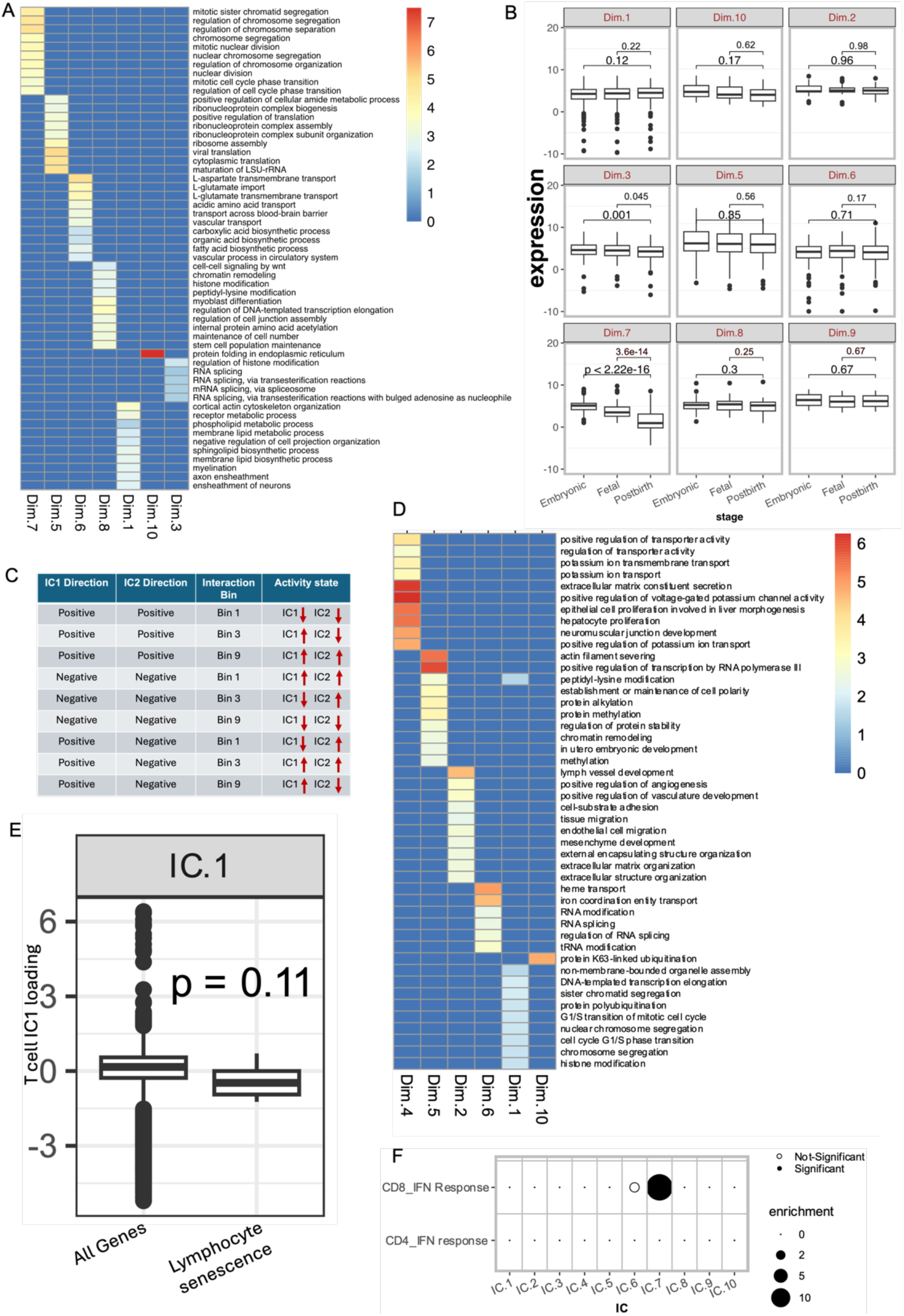
Identification of glioma stem cells and their interactions with other cell types in TME. **A.** A heatmap showing the GO terms enriched across the negative signature genes of the Malignant cell ICs. The cell colors show the odds ratio of enrichment for each significant GO term in at least one of the comparisons. **B** A boxplot showing the expression (in log TPM units) of negative signature genes of various malignant cell ICs during the embryonic development of human brain. P-values from Wilcoxon’s rank sum test are shown. **C** A schematic showing the activity state of positive and negative signature genes of interacting ICs across different activity bins. Arrows indicate upregulation (upwards arrow) and downregulation (downwards arrow). **D.** A heatmap showing the GO terms enriched across the negative signature genes of the T cell ICs. The cell colors show the odds ratio of enrichment for each significant GO term in at least one of the comparisons. **E** A box plot showing the contribution of lymphocyte specific senescence markers to IC1 of malignant cells. P-value from Wilcoxon’s rank sum test is shown. **F.** A dot plot showing the odds ratio for the overlap between negative signature genes of T cells IC7 and interferon response signaling in CD8 and CD4 T cells. The size of dots is proportional to the odds ratio derived from Fisher’s exact test, and p-values were adjusted for multiple comparisons across all the curated T cell state markers provided by Yanshuo et al.

Another interaction involving malignant cell IC7 (i.e. GSCs) occurred with T cell IC7 in Bin 9 associated with worse survival. Negative signatures of IC7 in T cells significantly overlap the markers of interferon response (Figure S2F, method). Since interaction in bin 9 involves the downregulation of negative signature genes of both interacting ICs, this interaction suggests that reduction in proliferative malignant cells when alongside the down regulated interferon signaling in T cells is associated with worse survival. We focused on the genes that underlie this pro-tumor interaction between downregulated stemness and T-cell interferon signaling by directly assessing the effect of their simultaneous downregulation on patient survival using the same statistical framework as the one employed to detect IC interactions (Methods). This analysis resulted in 129 significant pairwise gene combinations with almost all (128/129) having pro-tumor effect upon their simultaneous downregulation (Figure 3G). Among these, bona fide interferon response genes IFIT1 and IFIT3 in T cells exhibited a significant pro-tumor interaction with PTN and FBXO5 genes in malignant cells, respectively. In particular, PTN gene, which encodes for a secreted growth factor and a cytokine with potential immune-regulatory function (Sorrelle, Dominguez, and Brekken 2017). The interaction between PTN and IFIT1 implies that downregulation of PTN in malignant cells may suppress interferon signaling in T cells and the associated cytotoxic response (von Locquenghien et al.). Together, these results indicate a functional crosstalk and synergistic effects of activity of GSCs with distinct T cell states and gene expression programs associated with survival.

### IDH-mut Astrocytoma and Oligodendrogliomas exhibit differential CSI activities

Past research has identified two major subtypes of IDH-mut gliomas. Patients with IDH-mut Oligodendrogliomas (IDH-O) carry a co-deletion of chromosomal arms 1p/19q and generally have better prognosis than IDH-mut Astrocytomas (IDH-A). Furthermore, IDH-A and IDH-O tumors differ substantially in their TME composition (Venteicher et al. 2017). Therefore, we investigated if the detected CSIs had differential effects in these two tumor types. Towards this, for each interaction, we quantified a metric called ‘interaction penetrance’ (Methods) defined as the proportion of patients in which the interaction was active (for instance, the proportion of simultaneously downregulated patients for an interaction in Bin 1 and so on). In the TCGA cohort, we observed a significantly higher CSI penetrance (Methods) in IDH-A tumors compared with the IDH-O tumors (Figure 4A). To further assess potentially disparate roles of the CSIs in IDH-A and IDH-O tumors, we identified significantly differentially active CSIs between the two tumor types using Fisher’s test (Methods). We observed that CSIs dominant in IDH-A tumors were predominantly associated with anti-tumor activity (Figure 4 B, C; Fisher Odds = 2.26, one tailed p-value = 0.078). Since IDH-A tumors generally have poorer prognosis as compared to IDH-O tumors (van der Vaart et al. 2024), the enrichment of anti-tumor CSIs in IDH-A tumor is surprising. However, to further test the validity of pro-tumor and anti-tumor CSIs, separately for IDH-A and IDH-O cohorts, we quantified a metric called interaction load, defined as the total number of interactions active in each patient (Methods). Using interaction load of pro- and anti-tumor interactions, we partition the samples into those that had predominantly pro-tumor CSIs active and those that had predominantly anti-tumor CSIs (Methods; Figure 4D, E), and analyzed the survival patterns of these four groups of patients. We observed that for both IDH-A and IDH-O cohorts, patients dominated by anti-tumor interactions exhibit significantly better survival (Figure 4F, G). Thus, accounting for the activity of specific types of CSIs in a tumor refines the current understanding of the prognosis of the two subtypes of IDH-mut gliomas.

**Figure 4.**
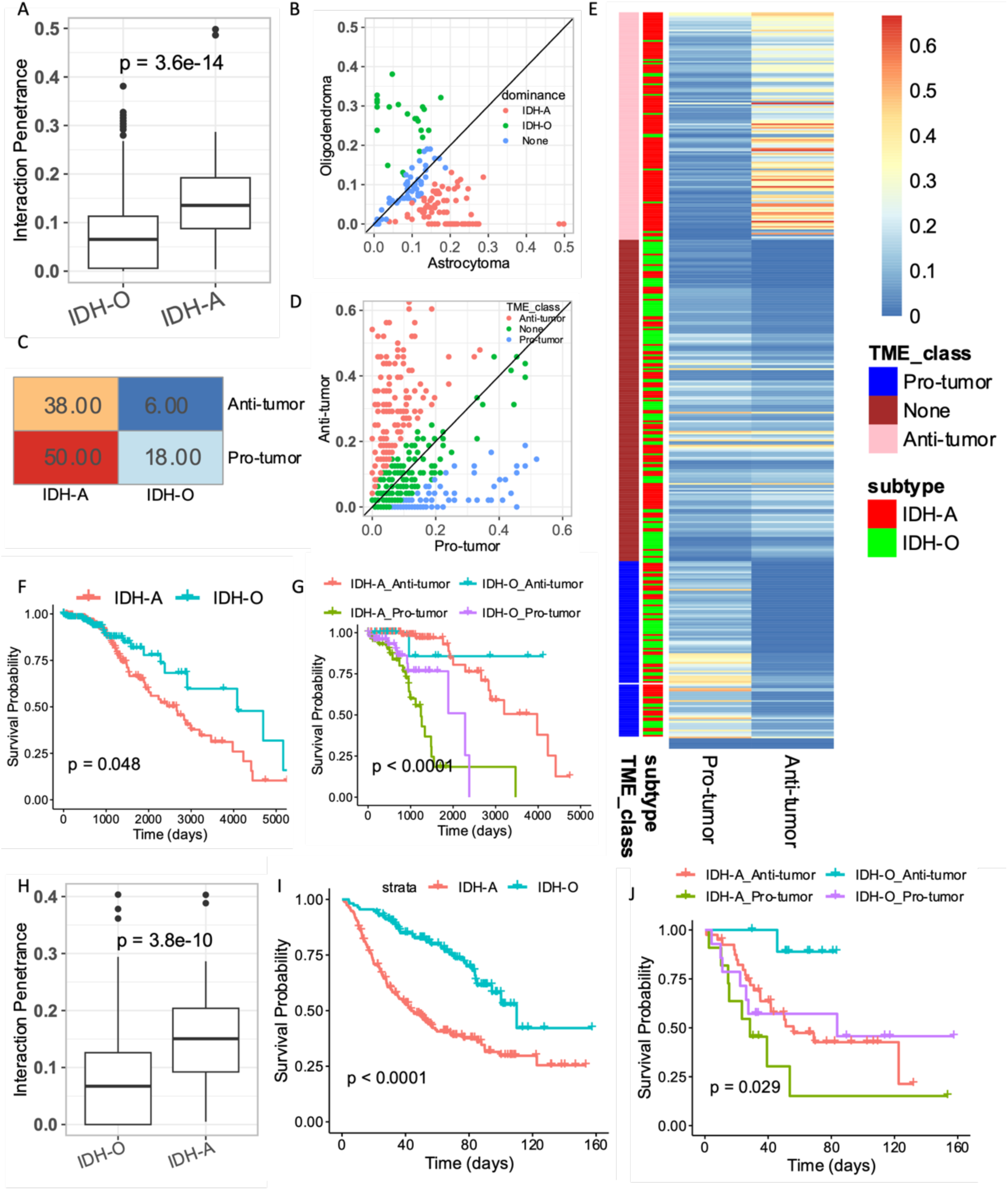
Analysis of CSIN in IDH-A and IDH-O tumors. **A** Boxplot showing the penetrance (i.e. proportion of patients where an interaction is active) of interactions among the IDH-A and IDH-O tumors in TCGA cohort. P-value from Wilcoxon’s rank sum test is shown. **B** Scatter plot showing the penetrance of each interaction in IDH-A and IDH-O in TCGA cohort. Each dot represents an interaction. The interactions enriched among the IDH-A are colored as red and the those enriched among IDH-O are colored blue. **C** Heatmap showing the numbers of dominant pro- and anti-tumor interactions active among the IDH-A and IDH-O subtypes in TCGA cohort. **D** Scatter plot showing the interaction load (i.e. the proportion of interactions active in a specific sample) for the pro- and anti-tumor interactions in TCGA cohort. The samples where pro-tumor interactions are dominant are colored as blue and samples where anti-tumor interactions are dominant are colored as red. These two groups of samples are used to define the dominant TME subtype. **E** Heatmap showing the interaction load of pro- and anti-tumor interactions for the samples with defined TME subtypes in panel “D” in TCGA cohort. **F** Kaplan Meier survival curves showing the overall survival probability of IDH-A and IDH-O in TCGA cohort. P-value from log-rank test is shown. **G** Kaplan Meier survival curves showing the overall survival probability of IDH-A and IDH-O stratified based on the dominant TME class defined in D in TCGA cohort. Patients with IDH-A tumors, despite the poor overall survival, show improved prognosis when dominated by anti-tumor interactions in the TME. **H** Boxplot showing the penetrance (i.e. proportion of patients where an interaction is active) of interactions among the IDH-A and IDH-O glioma subtypes in CGGA cohort. **F** Kaplan Meier survival curves showing the overall survival probability of IDH-A and IDH-O in CGGA cohort. **G** Kaplan Meier survival curves showing the overall survival probability of IDH-A and IDH-O stratified based on the dominant TME class in CGGA cohort.

We validated the observations made in the TCGA cohorts using an additional independent cohort of IDH-mut glioma patients from CGGA cohort (Methods). For this analysis, we considered the interactions that had similar direction of hazard in TCGA and CGGA (i.e. 103 /160 interactions) and scored the samples in CGGA for their joint activity (Methods). We observed that, consistent with TCGA cohort, stratifying the patients with IDH-A tumors based on their dominant TME subtype revealed a significantly improved survival of for the cases where TME is dominated by the anti-tumor interactions (Figure 4I, J). Taken together, these observations point towards the impact of differential interactions in TME on the overall survival of IDH-A and IDH-O which are not necessarily explained by co-deletion of 1p/10q chromosomal arms.

### Ligand-receptor interactions are likely to mediate CSIs

Intercellular communication in the tissue microenvironment is mediated, in substantial part, via ligands and receptors (Armingol et al. 2021). Here we aimed to assess the extent to which the detected CSIs are mediated *via* ligand-receptor (LR) interactions. Toward this, first we assessed whether the degree of interaction between a pair of cell types in the CSIN correlates with the number of known complementary LR pairs expressed by those cell types (Methods). Interestingly, we observed (Figure 5A and Supplementary Figure S3A) that the cell types which interacted more frequently in the CSIN were more enriched for complementary LR pairs among their ICs. Specifically, a total of 20% of the interactions in the CSIN (32 / 160) involved complementary LR pairs among their interacting components, comprising a total of 69 unique ligand-receptor interactions among the 7 cell types (Figure 5B).

**Figure 5.**
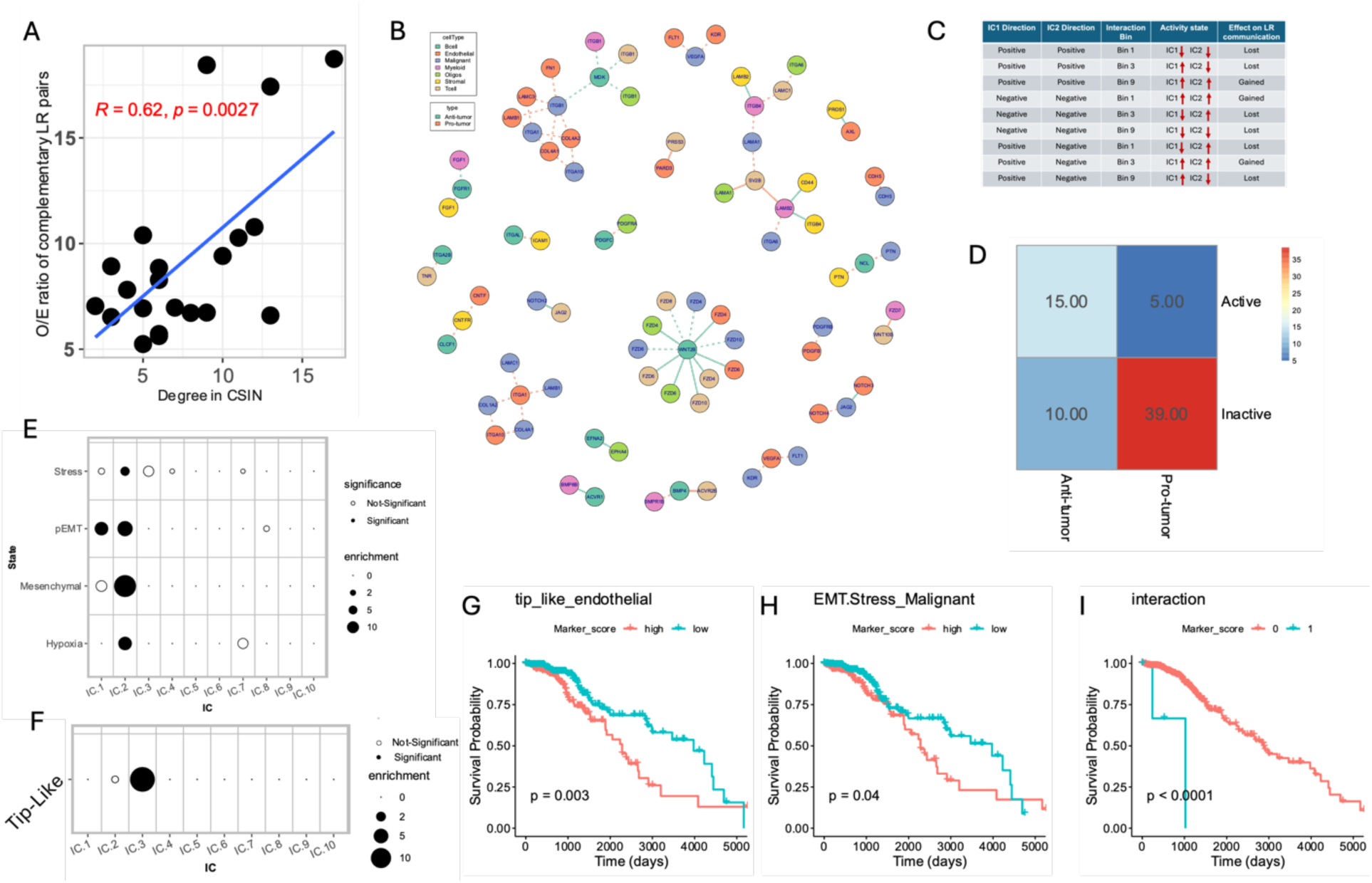
Analysis of ligand-receptor mediated cell state interactions. **A.** Scatter plot showing the degree of interactions between cell type pairs calculated from CSIN and observed by expected ratio of the complementary ligand receptor pairs present across all IC pairs between each cell type. Spearman’s correlation coefficient and corresponding p-value is shown in the plot. **B.** A network plot showing the known ligand receptor pairs that mapped to the interactions in CSIN. The text on each node is the gene name for the ligand/receptor and color represent the corresponding cell type in the CSIN. Color of edges indicate the pro-tumor (red) or anti-tumor (green) effect of the interactions and the shape of edge indicates the state of ligand-receptor interactions, i.e., activated (solid line) or inactivated (dotted line). **C.** A schematic table to explain that the presence of complementary ligand/receptor pairs among the negative or positive signature genes of ICs affects their activity across different interaction bins. The direction of arrows shows the status of positive and negative signature genes of their corresponding ICs across different interaction bins. A given LR interaction is inactivated if one or both partners are downregulated. **D.** A heatmap showing the number of activated and inactivated ligand receptor pairs having either pro- or anti-tumor interactions in the CSIN. **E.** A dot plot showing the odds ratio for the overlap between the positive signature genes of various Malignant cell ICs and various cancer cell states. **F.** A dot plot showing the odds ratio for the overlap between the negative signature genes of various T cell ICs and the markers of tip-like endothelial cells. In both **E** & **F**, the size of dots is proportional to the odds ratio derived from Fisher’s test and solid points indicate the comparisons where the FDR corrected p-value < 0.20. **G-I.** Kaplan Meier’s plot showing the survival probability of the IDH-mutant glioma patients stratified based on the activity of tip-like cells (i.e. IC3 of endothelial cells) in **G,** Hypoxic malignant cells undergoing EMT (i.e. IC 2 of the malignant cells) in **H**, and the interaction between these two in **I**.

We sought to identify which ligand-receptor interactions were activated or lost in the CSIN. Since a ligand and its receptors can be a member of either the positive or negative signature genes of the ICs, we extracted the joint activity state of the ligand and receptors in the CSIN by accounting for the sign of their corresponding IC and their interaction bin (Methods, Figure 5C). We identified a total of 20/69 activated ligand-receptor pairs, 15 (75%) of which were the part of anti-tumor interactions in CSIN. On the other hand, 49/69 identified ligand-receptor pairs were inactivated, 39 (80%) of which were part of pro-tumor interactions in CSIN (Figure 5D).

We next focused specifically on the ligand-receptor pairs whose deactivation is associated with pro-tumor effects. A majority (25/39) of these ligand-receptor pairs were present among the negative signature genes of 3^rd^ IC of endothelial cells and positive signature genes of 2^nd^ IC of the malignant cells and the interaction occurred in Bin 9. To understand the functional significance of this interaction, we overlapped their corresponding signature genes with known states of endothelial and malignant cells. Interestingly, we observed that the positive signature genes of 2^nd^ IC from malignant cells involved in this interaction mapped to multiple markers including hypoxia, stress, partial EMT as well as mesenchymal cell state (Figure 5E). Hypoxia induces the stabilization of hypoxia inducible factor (HIF) and has pleiotropic effects on the cell physiology including the activation of epithelial mesenchymal transition (Hapke and Haake 2020). Interestingly, the negative signature genes of the 3^rd^ IC of endothelial cells significantly resembled the markers of tip-like endothelial cells (Figure 5F, Odds ratio ~ 16 & FDR < 3.0e-06) which is a state of endothelial cells involved in the angiogenesis (del Toro et al. 2010). Angiogenesis is typically associated with worse prognosis, and indeed, we confirmed in our data that the 3^rd^ IC of endothelial cells had a negative hazard ratio implying the pro-tumor role of its negative signature genes (Figure 5G). Thus, while down-regulation of tip-like cells is associated with improved prognosis, our results suggest that it has an opposite (pro-tumor) effect, in conjunction with hypoxic malignant cells (Figure 5I), rooted in the loss of the LR-mediated communication between these two cell states (Figure 5B).

Another interaction, which occurred in 3^rd^ IC of the malignant cells and 2^nd^ IC of the T cells occurred in Bin 1 and was associated with better prognosis. These components contained NOTCH2 and JAG2 respectively among the negative signature genes of the ICs of the malignant and T cells, implying their simultaneous activation. Consistent with the role of NOTCH2-JAG2 in the adhesion of cells (Murata et al. 2014), we observed that negative signature genes of the 2^nd^ IC of T cells significantly resembled the markers of T cell adhesion (Figure S3B, methods). Therefore, this interaction indicates that attachment of T cells to malignant cells via JAG2-NOTCH interaction might be an important factor in T cell mediated anti-cancer immunity and consequently associated with the improved patient prognosis (Kelliher and Roderick 2018).

**Figure S3.**
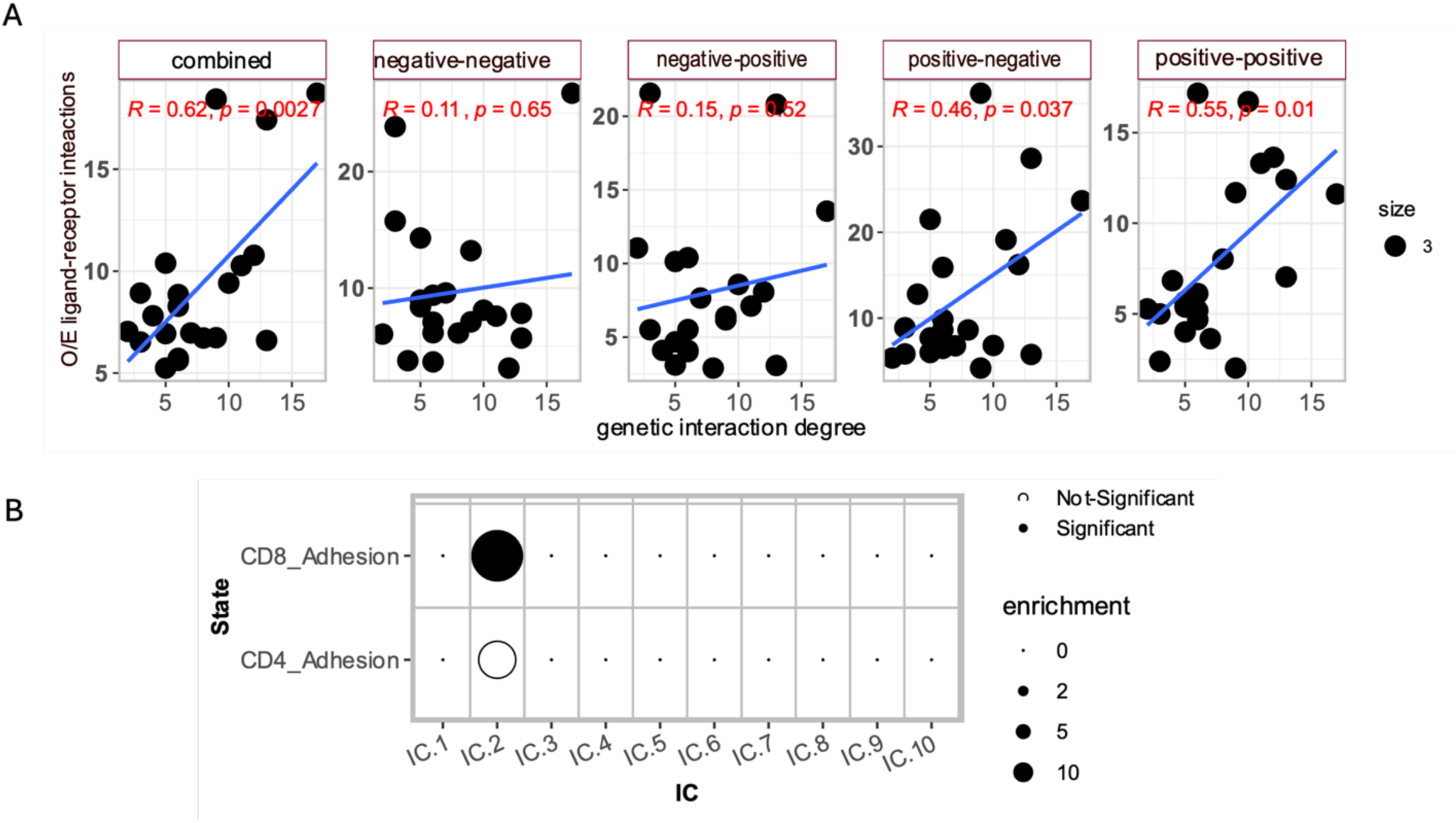
Analysis of ligand-receptor mediated cell state interactions. **A.** Scatter plot between the degree of interactions between cell type pairs calculated from CSIN and observed by expected (O/E) ratio of the complementary ligand receptor pairs present among the signature genes of all IC pairs between two cell types. We perform this calculation by considering all four possible combinations of the positive and negative signature genes of ICs as indicated at the top of each plot. Spearman’s correlation coefficient and corresponding p-value is shown in the plot. **B.** A dot plot showing the odds ratio for the overlap between the negative signature genes of various T cell ICs and markers of T cell adhesion. Size of dots is proportional to the odds ratio derived from Fisher’s test and solid points indicate the comparisons where the FDR corrected p-value < 0.20.

### CSIs are associated with therapy response and evolve during tumor relapse

Here, we assessed whether the identified interactions in the TME contribute to therapeutic resistance and how they change in relapsed tumors. Toward this, we first tested if the pro- and anti-tumor interactions were associated with response to immunotherapy and targeted inhibition of mutant IDH. We performed this analysis on a robust subset of interactions in CSIN consisting of 24 interactions (Methods, Supplementary File). We obtained a pre-treatment transcriptomic dataset from 29 patients undergoing neoadjuvant anti-PD1 immune checkpoint blockade therapy (Methods). We deconvolved this dataset using CODEFACS and in each sample, projected each cell type onto the ICA space established in the TCGA dataset (Methods), enabling us to determine the activity status of each cell type-specific IC in each sample. We calculated the penetrance of pro- and anti-tumor interactions in these patients (Methods) and, encouragingly, we observed that among the resistant patients, the penetrance of pro-tumor interactions was significantly higher than responders (Figure 6A). On the other hand, penetrance of the anti-tumor interactions was substantially higher in responders with marginally significant p-values, possibly attributable to the smaller sample size (N = 4 for anti-tumor interactions) (Figure 6A). These results indicate that cell state interactions might be crucial for mounting successful immune responses intended to be achieved by immune checkpoint blockade therapies.

**Figure 6.**
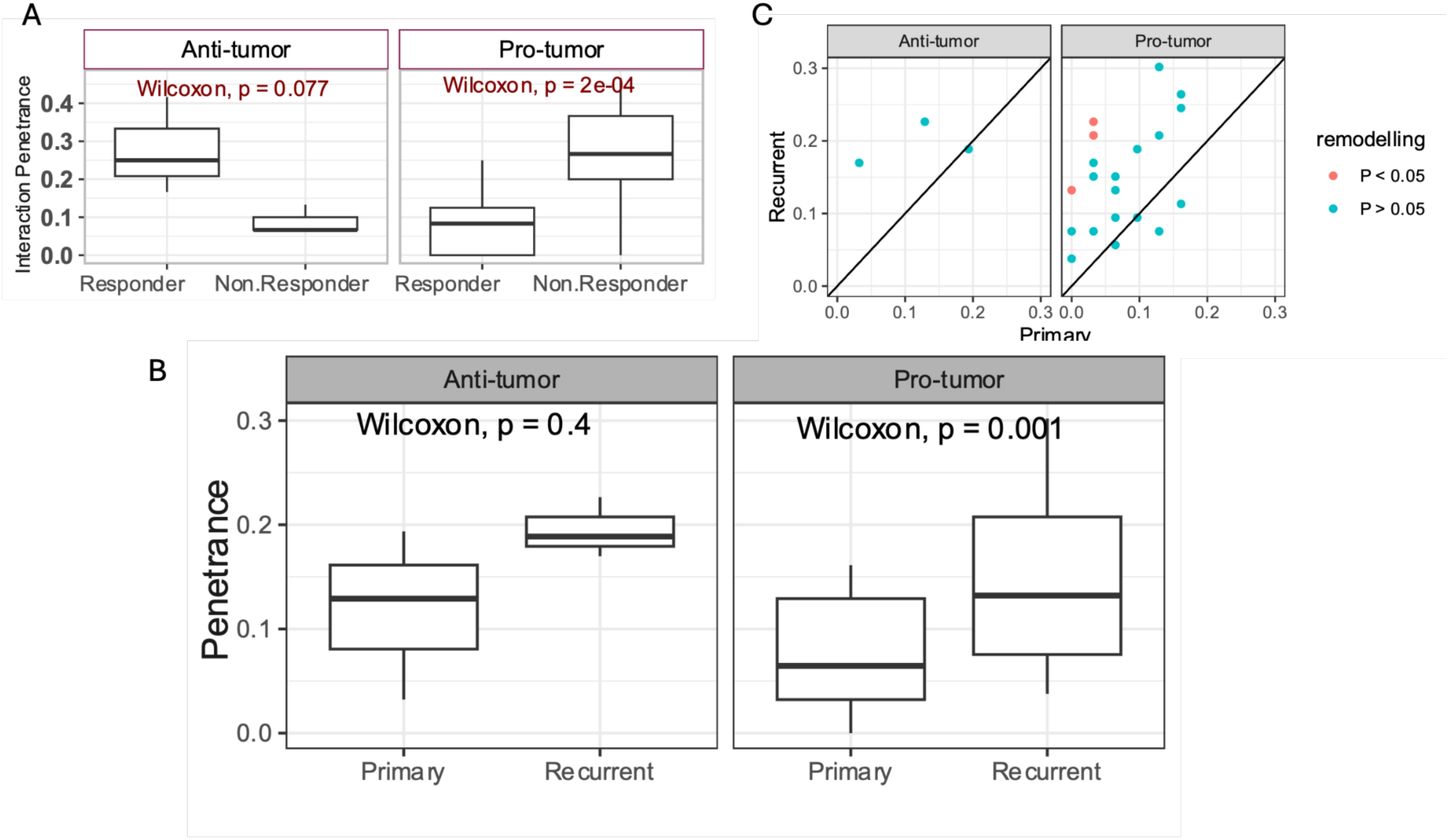
Association of CSIN with therapy response. **A.** Boxplots showing the distributions of penetrance of pro- and anti-tumor interactions in the CSIN among the responding and non-responding glioma patients that were administered immune checkpoint blockade therapy. **B.** Boxplots showing the distributions of penetrance of pro- and anti-tumor interactions in the CSIN among the primary and recurrent glioma patients. **C.** A scatter plot with the same data as in ‘b’ where each dot represents the penetrance of each pro- or anti-tumor interactions among the primary (x-axis) and recurrent tumors (y-axis). The interactions whose penetrance changed significantly (i.e. nominal p-value < 0.05) between primary and recurrent patients are highlighted by red colors.

Together, these results suggest that pre-existing interactions among the various cell states in the TME can significantly influence the outcome of targeted therapies. We further investigated how the interactions evolve as the tumor relapses after initial shrinkage following the standard of care therapies. For this, we obtained the bulk transcriptomic data from paired primary and recurrent tumor biopsies from a cohort of 25 IDH-mutant glioma patients collected by GLASS consortium (Methods). As above we generated the cell type-specific gene expression profiles using CODEFACS and projected them onto the ICA space established in the TCGA dataset (Methods). We hypothesized that following tumor relapse, there should be an increase in the number of pro-tumor cell state interactions as compared to the primary tumors. Directly comparing the interaction loads of the primary and recurrent biopsies from the same patients, we observed a significantly greater pro-tumor interaction load in the recurrent tumors as compared to the primary cases (Supplementary Figure S4A, B). We also observed a higher interaction load of anti-tumor interactions in recurrent biopsies, though not statistically significant (Supplementary Figure S4A). To identify the specific interactions underlying this trend, we tested for the interactions whose penetrance among the recurrent tumors was significantly enriched relative to the primary cases (Methods). Consistent with interaction load, we observed that penetrance of pro-tumor interactions was also significantly higher in recurrent tumors compared with primary cases (Figure 6B, C). At a nominal p-value cutoff of 0.05, we identified three interactions (Figure 6C) and all three had pro-tumor survival roles. Two of these interactions involved stromal cells which were recently proposed to stimulate glioma progression by regulating the tumor microenvironment (Cai et al. 2021). These results suggest that the pro-tumor interactions in the TME might mediate and get selected for in patients with relapsed tumors.

**Figure S4.**
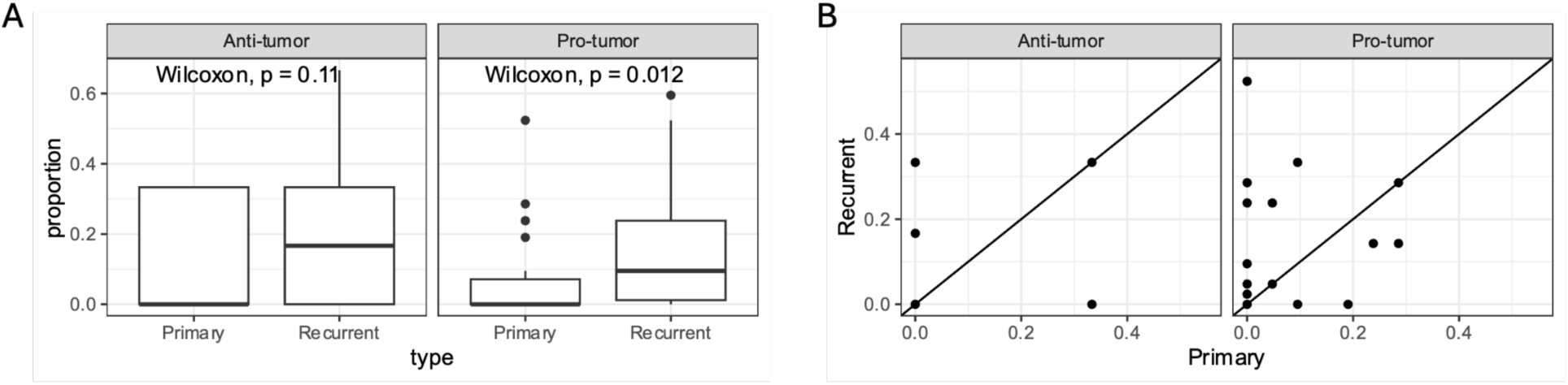
Association of CSIN with therapy response. **A.** Boxplots showing the distributions of interaction load for pro- and anti-tumor interactions among the primary and recurrent glioma patients. **B.** A scatter plot with the same data as in ‘B’. Each dot represents a paired value of interaction load derived from primary (x-axis) and recurrent (y-axis) tumors from the same patient.

### CSIs exhibit tumor stage-specific associations with somatic mutations

Somatic mutations in cancer cells can affect emergence of unique cell states within the TME (Ramirez et al. 2024), and shape the composition and functionality of the TME (Mansouri et al. 2022). Therefore, we explored the extent to which our inferred cellular states and their interactions are associated with the mutational events in cancer patients. Towards this, we first identified the ICs of 7 cell types which were significantly associated with specific non-synonymous somatic mutational events in patients of the TCGA cohort using a logistic regression approach (Methods). Screening through mutational status of 139 genes which were frequently mutated (defined as > 1% of the samples) and 70 ICs from 7 cell types, we observed a total of 135 significant associations (out of 9730 possible combination) between the activity level of the cell type ICs and mutational status of 9 genes, with mutations in TP53, ATRX, and CIC showing most frequent associations. (Supplementary Figure S5A, S6B).

These mutations were associated with 36 distinct ICs from 7 cell types, predominantly with Endothelial cells and Stromal cells (Supplementary Figure S5A). These observations hint at the influence of somatic mutations on the composition of the cell states in the tumor microenvironment.

**Figure S5.**
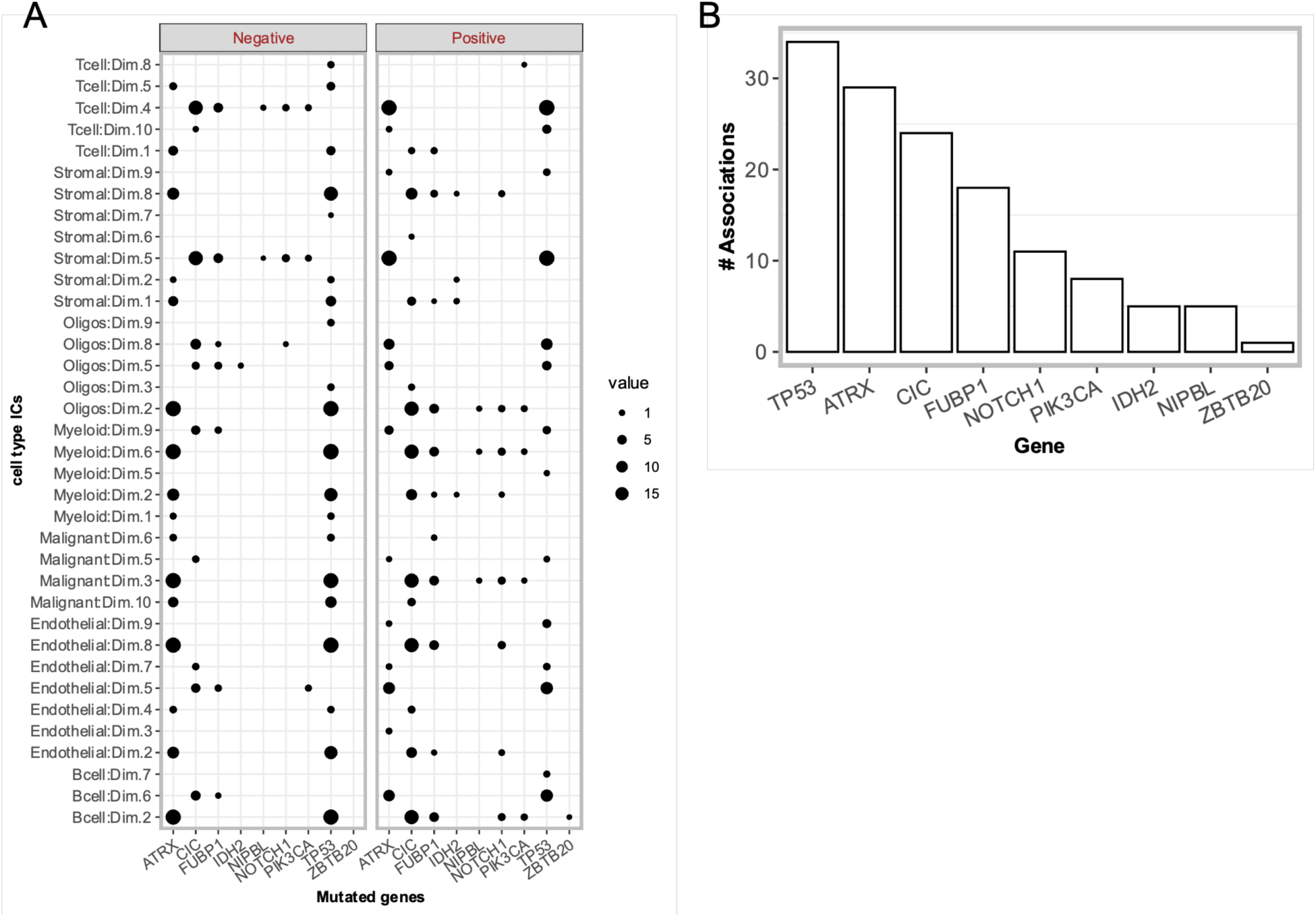
Association between somatic mutations and CSIN. **A.** A dot plot showing the odds ratio for the overlap between IDH-mutant glioma patients having specific nonsynonymous somatic mutations (x-axis) and high activity score (~top 33%) of ICs (y-axis). Size of dots is proportional to the odds ratio derived from Fisher’s test and solid points indicate the comparisons where the FDR < 0.20. Only the ICs and genes with at least 1 significant association (i.e. 91 associations) are shown. **B.** A bar plot showing the number of significantly associated ICs for each significant gene in **A**

We next investigated, beyond the ICs, if the detected IC interactions were also associated with the somatic mutations. Towards this, we first plotted the total mutational burden (Methods) against the total interaction load (as defined above) across the IDH-mut glioma samples. We observed a significant, albeit small, correlation between mutational load and interaction load for pro-tumor as well as anti-tumor interactions (Figure 7A), indicating the potential influence of cancer somatic mutations on the cell-cell interactions in the TME.

**Figure 7.**
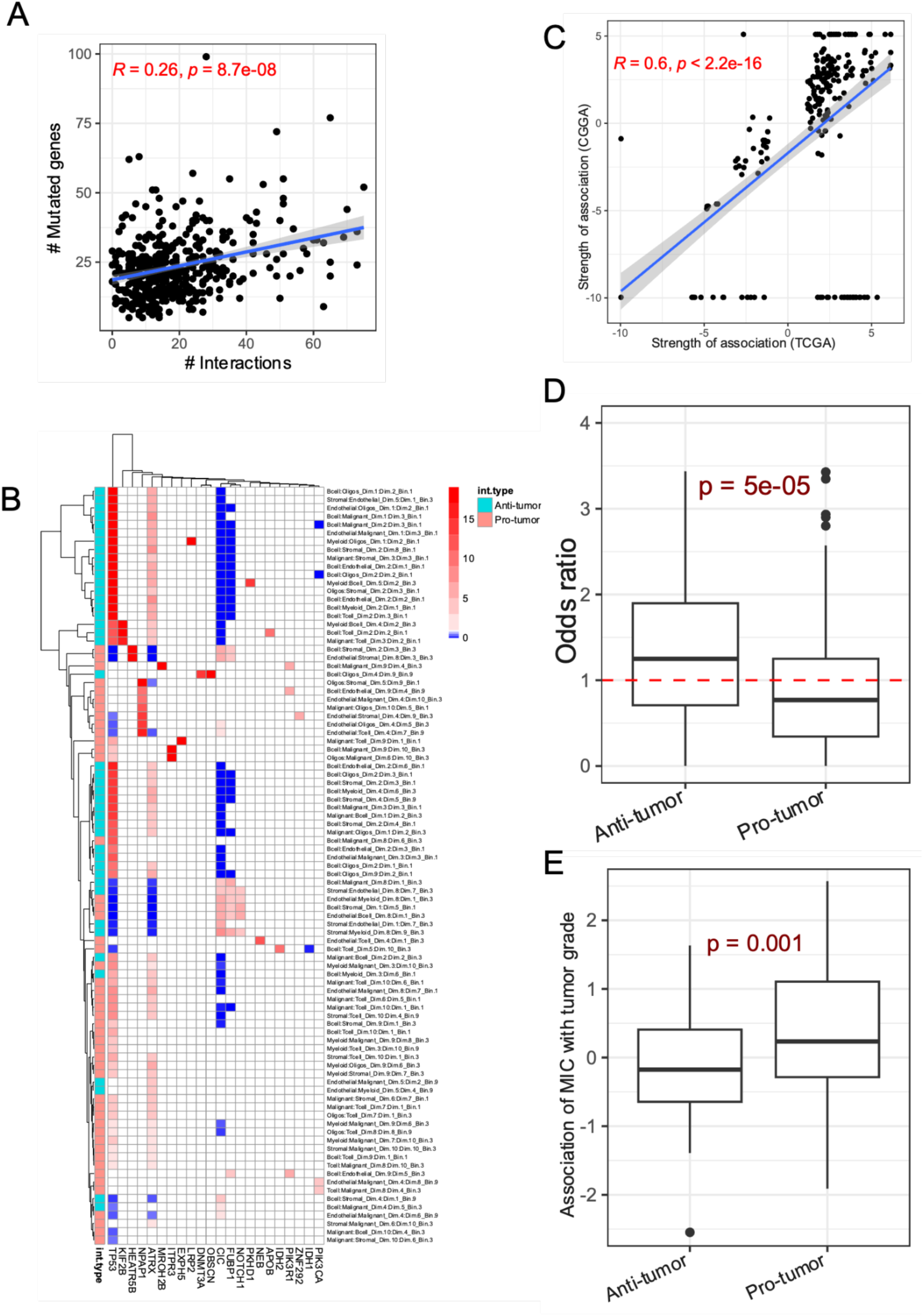
Association between somatic mutations and CSIN. **A.** Scatter plot between the interaction load (x-axis) and non-synonymous mutation load (y-axis) in the TCGA samples. The Spearman’s correlation and p-value are shown. **B.** A heatmap showing the strength of significant associations between specific interactions in CSIN and the nonsynonymous somatic mutations in the frequently mutated genes in IDH-mutant gliomas. Cell colors code the odds ratio of overlap between the mutated samples for each gene (x-axis) and samples where specific interaction (y-axis) was active. Only the interactions with at least 1 significant association (i.e. 91 interactions) are shown. The cases where association between a gene and interactions was not significant are colored as white. **C. A** scatterplot showing the correlation between the strength of mutation ~ interaction association (i.e. odds ratio) in TCGA and CGGA cohort of IDH-mutant glioma patients. The Spearman’s correlation and p-value are shown. **D.** Boxplot showing the enrichments (odds-ratio) of co-occurrence between somatic mutations and interactions in lower grade samples relative to higher grade for pro- and anti-tumor interactions. Dotted line shows the baseline expectation value under the null hypothesis that co-occurrence between interactions and mutations is independent of the tumor grade. **E.** Boxplot showing the estimate of the strength of association between mutation ~ interaction co-occurrence (MIC) with tumor grade for pro- and anti-tumor interactions. In both **D** and **E**, the p-values derived from Wilcoxon’s rank sum test are shown.

To identify the specific genes whose mutations are significantly associated with interactions in CSIN, we assessed the significance of the co-occurrence between cancer somatic mutations and interactions in the CSIN across the patients using a Fisher’s test (Methods). We observed a total of 266 instances of significant associations (FDR < 0.2) involving 22 mutated genes and 91 interactions from the CSIN (Figure 7B). A majority (~64%) of these significant associations involved the enrichment (as opposed to depletion) of interactions among the mutated patients, consistent with the overall positive correlation between mutation burden and interaction load (Figure 7A). We assessed the robustness of these associations between cancer somatic mutations and cell state interactions by performing a similar analysis on the independent CGGA cohort of IDH mutant patients (Methods). We observed the odds ratio of mutation-CSI association between these two cohorts to be highly correlated, highlighting the reproducibility of the relationship between somatic mutations and CSIs (Figure 7C).

Quite surprisingly, these 91 mutation-associated interactions spanned a majority (43 / 48) of the anti-tumor interactions in the CSIN despite their overall depletion in the CSIN (a total of 48 anti-tumor vs. 112 pro-tumor interactions in CSIN). In other words, somatic mutations are more likely to be associated with an anti-tumor interaction than with a pro-tumor interaction, raising the possibility that the tumor microenvironment may be eliciting a homeostatic response against somatic mutations in cancer to restore normal physiology. We reasoned that if true this would imply a greater association between anti-tumor interactions and the specific somatic mutations in earlier stages of cancer as compared to the later stages. To test this for a specific mutation and CSI, focusing on samples where the CSI is active, we compared their early (grade I & II) versus late (grades III & IV) split for mutated and the wildtype samples as an odds ratio, quantifying the tendency for the mutation and CSI to co-occur in early stages (Methods). We observed that for anti-tumor interactions, the early:late ratio was significantly higher for mutated samples as compared to the unmutated samples (Figure 7D). The pro-tumor interactions, on the other hand, showed a reverse trend. These results are consistent with our hypothesis that anti-tumor CSIs may, in part, be an early homeostatic response to the oncogenic somatic mutations, and as the tumor progresses to more advanced stages the TME is reprogrammed to support tumor progression via pro-tumor CSIs. We further strengthened this finding through an alternative approach. We defined a binary variable called mutation interaction co-occurrence, which is “1” for a sample where the two co-occur and “0” otherwise (MIC, Methods), and modeled the relationship between tumor grade and MIC while controlling for the effect of mutation and interaction alone (Method). Consistently, we observed MIC was negatively associated with tumor grade for anti-tumor interactions and positively associated with pro-tumor interactions (Figure 7E), suggesting that early on, anti-tumor interactions are enriched in mutated samples relative to the wildtype samples, possibly to restore normal physiology of the underlying tissue.

## Discussion

Genes evolve and function in the context of the genetic and regulatory networks (Phillips 2008), a phenomenon termed epistasis in molecular genetics. The knowledge of such interactions has implications in precision oncology and combination therapy (Szczurek, Misra, and Vingron 2013). One special case of gene interactions is synthetic lethality, where the joint, but not individual, inactivity of two genes results in reduced cell viability, exemplified by the discovery and FDA approval of PARP-inhibitors which are particularly effective in BRCA deficient cancers because of the synthetic lethal interactions between these genes (Helleday 2011). These ideas have fueled the development of several data driven methods to identify clinically relevant interactions between genes (Jerby-Arnon et al. 2014; Lee et al. 2018; Magen et al. 2019; Matlak and Szczurek 2017).

In this work, we extend the concept of functional interactions between genes to those between cellular states across the cell types in the TME. A myriad of cell types are present in TME in addition to the malignant cells. While single cell and spatial transcriptomics datasets from cancer patients provide a robust means to understand the composition, heterogeneity and intercellular communication (Cheng et al. 2023), due to technical difficulty and cost, such data substantially lag behind the rich clinical information available in large cohorts of bulk transcriptomic data such as TCGA. In this work, we developed a novel computational pipeline that leverages the large cohorts of clinical bulk transcriptomics data to understand clinically relevant crosstalk in TME.

Applying our method to bulk transcriptomic and clinical data from TCGA IDH-mutant glioma cohort, we detected interactions involving multiple cellular states in the TME. The resulting cell state interaction network had a greater than two-fold enrichment for pro-tumor interactions relative to the anti-tumor interactions, suggesting that the TME is globally wired to promote tumor progression. The detected interactions were significantly reproducible in another independent cohort of IDH-mutant glioma patients ensuring the technical as well as biological robustness of the CSIN. Biologically meaningful nature of our approach is also underscored by the fact that we could recover multiple known states of various cell types present in the TME through our approach. For instance, through the ICA of deconvolved bulk transcriptomic data, we could identify multiple malignant cell states which have been previously proposed to exist in IDH-mutant gliomas including GSCs, Astrocytic-like and OPC-like. Further, the ICs derived from deconvolved cell types had a significantly consistent effect on patients’ survival between TCGA and CGGA, ensuring the technical robustness of our approach and enabling us to assess their effect on patient survival.

Single cell RNA sequencing, combined with known markers of various cell types, has helped characterize cell type compositions in a tissue. However, reliance on a handful of known markers cannot reveal subtle shifts in transcriptional states of various cell types. Latent space embedding of the scRNA-seq data using non-negative matrix factorization (Kotliar et al. 2019), or supervised learning approaches (Kunes et al. 2024) can provide a more resolved view of transcriptional states of various cell types in the tumor microenvironment. Given a resolved cell state composition, intercellular communication can be inferred based on the expression of complementary ligand-receptor pairs by distinct cell types (Armingol et al. 2021); however, our knowledge of functional ligand-receptor pairs is limited. The broader goal, however, is to understand how the cell state composition and intercellular interactions vary across tumors and whether those variations are linked to clinical features, to ultimately be modulated for therapeutic purposes. A previously developed method called Dialogue can bypass the need for a database of known ligand-receptor interactions to infer cell-communication by defining a concept of multiple cellular communities based on the canonical correlation analysis of cell type specific gene expression programs in scRNA-seq data (Jerby-Arnon and Regev 2022). However, it only captures the cell states or gene expression programs that co-vary between multiple cell types, unlike CSI-TME, which infers prognostic cell state combinations that do not necessarily co-vary across patients. The main challenge toward the broader goal stated above is insufficient clinical scRNA-data. CSI-TME overcomes these limitations by (i) deconvolving a large cohort of clinical bulk transcriptomic data, (ii) identifying cell states and their transcriptional programs, and (iii) explicitly identifying clinically relevant cell state pairs based on their joint activity across cell types in the TME. Previously, a non-negative matrix factorization (NMF) based approach, called EcoTyper, was developed to identify multicellular tumor ecosystems across cancer types in bulk, single cell, or spatially resolved gene expression data (Luca et al. 2021). Tumor ecotypes were defined based on the significant co-occurrence of NMF based cell states across multiple tumor types. However, in CSI-TME, we infer interactions between cell state pairs by directly modeling the survival data based on joint activity/inactivity of the pairs of cell state ICs, a key difference that enable us to identify prognostic cell-cell interactions

It is intriguing that despite the lack of immunotherapy response information in TCGA, the prognostic cell state interactions that we identified in TCGA were associated with clinical response to PD-L1 blockade immunotherapy in an independent cohort. Since the cytolytic activity of immune cells in TME contributes to the survival of patients, the cell state interactions associated with worse patient prognosis in TCGA might represent a microenvironment conducive for tumor growth with low cytotoxic activity. Patients with such a microenvironment might not benefit from the PD-L1 blockade which is intended at activating the suppressed immune cells. One could potentially use CSI-TME to prioritize ligand receptor interactions underlying pro-tumor microenvironment and/or resistance to immunotherapy. Further, CSI-TME models the overall survival of patients based on the joint activity of distinct cell state combinations. However, given sufficient cohort size, CSI-TME can be directly used to model other clinical variables of interest such as response to therapies, tumor relapse, or metastasis based on the activity of TME.

Lastly, we observed that the anti-tumor interactions in the TME were differentially enriched among the samples bearing oncogenic mutations in early stages relative to the non-mutant samples. On the other hand, in later stages of tumorigenesis, this balance shifts towards the pro-tumor interactions. These results indicate the potential protective role of the TME during initial stages of tumor which gradually acquires the tumor promoting characteristics.

We note that transcriptional deconvolution of bulk gene expression data is a rapidly evolving field (Im and Kim 2023) and in addition to the core model assumptions, the accuracy depends on several factors such as sequencing depth, cell type abundance, and inter-tumor heterogeneity (de Vries et al. 2020). Further, the deconvolution accuracy might be improved by incorporating additional data modalities such as DNA methylation (Im and Kim 2023). In this work, we used a recently published transcriptional deconvolution method called CODEFACS which uses several heuristics to maximize the number of genes that are deconvolved confidently with an option to exclude low confidence genes (Wang et al. 2022). The observed consistency between two different cohorts (i.e. TCGA and CGGA) based on the ICA of the deconvolved cell types (Figure 2B, C) and recovery of known cell states (Figure 3A) supports the accuracy and robustness of the approach.

Taken together, leveraging the large cohorts of bulk transcriptomic and clinical datasets, our work provides a data driven computational pipeline to discover a significantly reproducible set of prognostic cell state combinations, prioritize clinically targetable ligand-receptor interactions, and stratify patients for immunotherapy.

## Methods

### Bulk RNA seq datasets

We used publicly available bulk RNA-seq datasets from IDH-mutant glioma patients from The Cancer Genome Atlas (TCGA) and Chinese Glioma Genome Atlas (CGGA) (Zhao et al. 2021). Read count matrix for 425 IDH-mutant glioma patients from TCGA was downloaded from UCSC xena browser (Goldman et al. 2020) and was normalized to TPM (Transcripts Per Million) using a shiny R based COEX-seq package (https://github.com/kimsc77/COEX-seq). To ensure the robustness and reproducibility of our findings, an additional bulk RNA-seq dataset (ID CGGA693) comprising 325 IDH-mut samples was downloaded from Chinese Glioma Genome Atlas (CGGA) in TPM units (http://www.cgga.org.cn/) (Kim, Yu, and Cho 2018). Raw RNA-seq data for developing the human brain was obtained from (Cardoso-Moreira et al. 2019) and processed to generate gene level TPM as part of a previous publication (Singh et al. 2022). RNA-seq data for longitudinal biopsies from primary and recurrent IDH-mutant gliomas was sourced from GLASS (Glioma Longitudinal Analysis) consortium (http://www.synapse.org/glass) and consisted of 85 RNA-seq profiles comprising primary and recurrent tumors from 31 patients-seq data. RNA-seq data from pre-treatment biopsies from 29 pembrolizumab treated glioma patients was sourced from a previous publication (Cloughesy et al. 2019).

### Clinical for TCGA and CGGA

Clinical data including the overall survival, age, sex, tumor grade, and IDH-A/IDH-O classification of IDH-mut glioma patients in TCGA was downloaded from <u>PanCanAtlas</u> (Nawy 2018). Similar clinical data for IDH-mut patients in CGGA cohort was downloaded from their <u>online portal</u>.

### Single cell RNA-seq dataset

scRNA-seq dataset consisting of 60751 cells from 4 IDH-mutant tumors was obtained from a previous publication (Abdelfattah et al. 2022) (<u>GSE182109</u>1) and 9 IDH-mutant tumors from in-house samples at NCI. Read demultiplexing and alignment to the GRCh38 human reference genome was performed using the Cell Ranger Single Cell Software v2.0 (10X Genomics) with its mkfastq and count functions, respectively. Downstream analysis was performed on the filtered counts with cells having at least 200 genes and the percentage of mitochondrial unique molecular identifiers (UMI) counts less than 20%. Filtered cells were then merged, batch-corrected with Harmony (Korsunsky et al. 2019) and clustered using Seurat v3 package using 5000 most variable genes (based on the “vst” selection method) at 0.1 resolution. Malignant cells were identified using CONICsmat and CopyKat V0.1.0 with default parameters (Müller et al. 2018) (Gao et al. 2021) while the remaining nonmalignant cells were annotated by using cell type-specific markers using provided in Supplementary Table S1. Finally, all cells were classified into 7 major cell types, viz., Malignant, Myeloid, B cells, T cells, Endothelial cells, Oligodendrocytes, and Stromal cells.

### Pipeline for CSI-TME

To identify cell state interactions, CSI-TME employs the following three main steps.

#### Deconvolution

In the first step, CSI-TME employs a recently published transcriptional deconvolution tool CODEFACS (COnfident DEconvolution For All Cell Subset) to deconvolve bulk RNA-seq datasets. For a given set of cell types along with their transcriptional signatures derived from single cell RNA-seq data, CODEFACS estimates cell type-specific expression profiles in a bulk transcriptomic data. We first supplied the annotated scRNA-seq data to CIBERSORT (Chen et al. 2018) with default parameters to generate the signatures for seven cell types – B cells, T cells, Malignant cells, Myeloid cells, Oligodendrocytes, Stromal cells, Endothelial cells. Using CIBERSORT’s Impute Cell Fraction module with its S-mode batch-correction, these signatures were then used to estimate cell type fractions in each bulk transcriptomic sample in large glioma datasets from TCGA and CGGA. Finally, CODEFACS was applied to derive their cell type-specific gene expression profiles in each TCGA and CGGA sample. For down-stream analysis we only considered genes that had a CODEFACS estimated confidence score > 0.95.

#### Cell state identification using ICA

Independent Component Analysis (ICA) is a computational technique to separate a multivariate signal into additive, statistically independent components. Recently, ICA is being increasingly used to model the transcriptomic datasets to uncover hidden patterns in complex cancer omics datasets (Sompairac et al. 2019). CSI-TME uses ICA to identify distinct cell states or gene expression programs, independently for each of the seven cell types, based on their deconvolved gene expression profile across tumors. We performed ICA using the JADE algorithm implemented in the MineICA package (Biton 2024) in R. We arbitrarily choose to extract 10 independent components for each cell type, reasoning that it should be sufficient to capture the transcriptomic and functional heterogeneity in a cell type. Prior to ICA, the input gene expression profiles of cell types were log transformed and z-scored across the samples. From the output of MineICA, source matrix (sample by components) was used as the estimate of sample specific score of latent cell states or their gene expression programs.

#### Screening for prognostic cell state combinations

In the last step, CSI-TME screens for the pairs of ICs across different cell types whose joint activity or inactivity is significantly associated with the patient’s survival. For this purpose, we first partitioned the sample specific scores of each IC into three equal parts defining three activity bins, viz., low, medium, and high. An IC is considered ‘active’ in the ‘high’ portion, and ‘inactive’ in ‘low’ part; ‘medium’ part was excluded. Next, for each pair of ICs (say, IC1 and IC2) taken from two different cell types, we assigned each sample to one of the three joint activity combinations (Low-Low, Low-High or High-Low, and High-High); Low-High and High-Low are treated equivalently without loss of generality. Lastly, iterating through each pair of ICs in each of the three joint activity bins, we modeled the overall survival of patients using Cox-proportional hazard model based on joint activity of ICs while controlling for the effect of each individual ICs as described below:

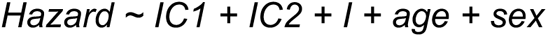

Where *IC1* and *IC2* are the sample-specific scores of the two *ICs*, and *I* code for the interactions, i.e., the joint activity bin of the *IC1* and *IC2*. Age and sex were used as covariates to control for additional demographic confounders. From this model, the hazard ratio and p-values for the interaction term *(I)* were considered and multiple testing correction based on Benjamin-Hochberg’s procedure was performed for each IC pair. The output of CSI-TME consisted of list of tuples (C1, IC1, C2, IC2, B, D) at an FDR threshold of 0.20, where IC1 and IC2 are the two ICs from cell type C1 and C2 respectively, and B represents their joint activity state and D represent the direction of prognostic association between hazard and interaction term, i.e, whether the joint activity state associated with better or worse survival.

Finally, CSI-TME performs internal cross-validation of the detected interactions by creating 10 bootstrapped sampling of the clinical data and repeating step 3 to detect significant interactions. CSI-TME only retains the interactions that had a consistent effect on patient survival with p-value < 0.05 in at least 70% of the trials.

### Projection of external datasets onto the ICA space of TCGA dataset

The CSIs detected in the discovery cohort of TCGA patients were validated and assessed in different contexts: (i) CSIs were validated in an independent CCGA bulk RNA-seq dataset; (ii) CSIs were evaluated in response to immunotherapy in Cloughsey et al. dataset (Cloughesy et al. 2019); and (iii) CSIs were evaluated in relapsed tumors from GLASS consortium (Barthel et al. 2019). In each of these three cases, to quantify the ICs discovered in the TCGA cohort, we first deconvolved the bulk RNA-seq dataset using the CODEFACS and signature genes of 7 cell types as before. Next, we projected the RNA-seq profiles of each purified cell type onto the latent space of independent components established in the TCGA dataset by performing the matrix multiplication of the loading matrix (i.e. mixing matrix) with the new input data to be projected (i.e. deconvolved gene expression profiles of the corresponding cell type).

### Statistics and data visualization

All statistical and computational analysis were carried out in the R programming language. The p-values for testing the significance of difference between two distributions were estimated using Wilcoxon’s rank sum test. Significance of overlap between different gene signature sets was computed using Fisher’s exact test. In case of multiple comparisons, p-values were adjusted for FDR using Benjamini-Hochberg’s method. We used a uniform FDR threshold < 0.20 for reporting significant associations in this study. We used the ggplot2 library in R to create most of the graphics in this work. The network layouts of the cell state interactions which were created using the combination of **network** and **igraph** package in R. Heatmaps were created using **pheatmap** library in R. Gene ontology analysis of the positive and negative signature genes of the ICs was performed using **clusterProfiler** package in R and significantly enriched terms (FDR < 0.20) were plotted as heatmaps.

### Signature genes of ICs

In ICA, the original data **X** with n features (genes) and m samples is modeled as **X**=**AS**. In this setup for k number of extracted components, **A** is the loading or mixing matrix consisting of n genes and k components. For each independent component, genes which were at least 2.5 standard deviations away from the mean column mean in **A** were taken as signature genes of the independent components. Since a priori we do not know which direction of the component is associated with a specific biological process, we extracted positive signatures (i.e. 2.5 standard deviation above the mean) and negative signature (i.e. 2.5 standard deviation below the mean) for each component separately.

### Scoring single cells using signature genes of ICs and assessment of clustering in PCA space

We used the signature genes of distinct ICs derived from each cell type and scored each individual cell of the corresponding cell type in single cell data using the *AddModuleSocre* function imported from the Seurat package in R (Stuart et al. 2019). Positively scoring cells indicates that a given signature gene set is enriched in those cells. To assess if the cells expressing the signature genes represent distinct cell states, we computed the Euclidean distances in PC space of scRNA-seq data among all pairs of positively scoring cells (i.e. within cluster) and compared it against the Euclidean distances between all pairs of enriched and non-enriched cells (i.e. between cluster). A significantly smaller distance (FDR < 0.20 & at least 10% shorter) among the positively scoring was taken to indicate clustering of IC signature genes in PC space.

### Marker genes of various cell states

Markers for recurrent cancer cell states were obtained from Berkley et al. (Barkley et al. 2022). Markers for T cell states were obtained from Yanshuo et al. (Chu et al. 2023). Markers for Endothelial cell states were obtained from Zhang et al. (J. Zhang et al. 2022). Markers for the lymphocytic senescence were obtained from Martyshkina et al. (Martyshkina et al. 2023). Markers for fibroblast senescence were obtained from Wechter et al. (Wechter et al. 2023). We used two different glioma-specific putative stemness signatures to assess the stemness features among the malignant cell ICs. PSS1 was defined based on the genes whose log-normalized gene expression (in TPM units) was significantly correlated (FDR < 0.2 & PCC > 0.5) with the log-normalized gene expression of the *IDH1* gene in the bulk transcriptomic data from TCGA. PSS1 consisted of 1140 genes in total. PSS2 was sourced from Venteicher et al. (Venteicher et al. 2017).

### Enrichment of known cell state markers among the ICs

To assess the extent and significance of overlap between the signature genes of ICs and known transcriptional or cell state markers, we computed the odds ratio and p-value using Fisher’s exact test. Whenever multiple comparisons were involved, we adjusted the resulting p-values with the Benjamini-Hochberg method.

### Calculation of the interaction penetrance and interaction load

For each interaction in CSIN, we assigned a binary status to each patient, indicating whether the interaction is active (assigned as 1) or inactive (assigned as 0). For instance, an interaction in bin.1 would imply that, for patients where two ICs are simultaneously downregulated, the interaction is marked as active (assigned a value of 1); otherwise, it is assigned a 0. This process results in a binary matrix, where the rows represent interactions, and the columns represent patients.

The penetrance of each interaction was calculated as the sum of the values in each row of this binarized matrix divided by the total number of columns (i.e. patients), representing the fraction of patients in which the interaction is active (i.e., has a status of 1). On the other hand, the interaction load measures how many interactions are active in each sample and is defined as the proportion of interactions that are active in each patient sample.

### Somatic mutation data

For somatic mutation data for TCGA IDH-mutant glioma patients, we downloaded the .maf files derived from whole exome sequencing from GDC using the TCGAbiolinks library in R (Colaprico et al. 2016). The resulting .maf files were converted to a gene by sample matrix with each entry denoting the number of non-synonymous somatic mutations. The somatic mutation data for CGGA IDH-mutant glioma patients was downloaded as the gene by sample matrix format from CGGA consortium website (http://www.cgga.org.cn/).

### Association of somatic mutations with ICs and CSIN

We first identified the genes that were recurrently mutated (at least 1 non-synonymous mutation in > 1% of the samples) in TCGA IDH-mut glioma patients, resulting in a total of 139 genes. Next, we divided each IC derived from the 7 cell types into three binned activity states (coded as 0,1,2 from low to high) using quantile binning. To investigate the association between somatic mutations and ICs, we iterated through all possible gene-IC pairs (i.e. 139 genes x 70 ICs) and used a binomial regression to model the binned activity level of ICs based on the somatic mutation status of each gene. We extracted the resulting regression coefficients and p-values from the model, adjusted for multiple comparisons using Benjamini-Hochberg’s method and retained the significant associations with FDR < 0.20. To investigate association between somatic mutations and cell state interactions, we screened through all possible genes-interaction pairs (139 genes * 160 interactions) and assessed the overlap between patients containing somatic mutation in each gene and patients where the interaction was active using Fisher’s exact test. The resulting odds and p-values were extracted and adjusted for multiple comparisons Benjamini-Hochberg’s method. Significant associations with FDR < 0.20 were retained.

### Test for differential association between mutation and interactions relative to tumor grade

Here we assess whether there is a greater enrichment of anti-tumor interactions among the mutated samples in early stages of tumor as compared to the later stages when tumor has already progressed. To investigate this, for each somatic mutation having significant association with one or more cell state interactions (Figure 7B), we first partitioned the samples into a mutant and wildtype relative to the specific somatic mutation, resulting in 22 such partitions corresponding to the mutational status of 22 genes. Next, for each of these 22 genes, we collected its significantly associated interactions, and for each interaction, and counted the number of samples where interactions were active in early stages (i.e. WHO grade II) and later stages of tumor (i.e. WHO grade III/IV). For each gene and each associated interaction, this exercise resulted in the following contingency table:

|  | Early Stage | Late Stage |
| --- | --- | --- |
| Mutant | c1 | c2 |
| Wildtype | c3 | c4 |

where c1, c2, c3, c4 are the counts of samples where an interaction is active (based on the joint activity status of the two ICs). For each mutant and wildtype rows, we calculated the relative proportion of interactions in early vs. late stages (i.e. r1 = c1/c2 for mutant samples and r2 = c3/c4 for wildtype samples). The odds of differential enrichment in early stages relative to late stages were then obtained by normalizing r1 against r2 as r1/r2. Finally, we plotted the odds for pro- and anti-tumor interactions as boxplots.

As an alternative, we directly assessed the association between mutation interaction co-occurrence (MIC) and tumor grade using logistic regression. For each somatic mutation and interaction (Figure 7B), MIC was defined as follows,

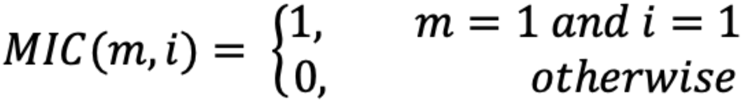

where *m* denotes the somatic mutation status of a given gene and *i* denotes the activity state of a given interaction across samples. MIC score of 1 implies that a given sample is both mutated as well as has the active interaction. The strength of association between MIC and tumor grade was then estimated by regressing tumor grade against the MIC, while controlling for the individual effect of *m, i* on tumor grade through a logistic regression model.

### Analysis of ligand-receptor interactions

Ligand-receptor pairs were sourced from cellchatdb database (Jin et al. 2021) consisting of 1986 interactions between 547 ligands and 466 receptors. For each cell type pair, we counted the number of complementary ligand receptor pairs among the signature genes across all combinations of their ICs. We accounted for the difference in the size of signature gene sets by dividing the observed count with the expected number of ligand-receptor pairs. For two ICs having with *n* and *m* number of signature genes, and *c* number of complementary ligand-receptor pairs, we normalized the count *c* as to derive observed by expected (O/E) ratio as follows:

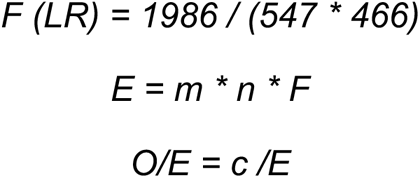

Where *F (LR)* is the global density of ligand-receptor connections among a total of 547 ligands and 466 receptors in cellchatdb database. *E* is the expected number of connections between two sets of *m* and *n* genes given a global density value of *F.* Finally, *O/E* denotes the observed count of *c* connections between *m* and *n* genes normalized by the expected value. For each cell type pair, we calculated the spearman’s correlation between O/E ratio against the number of interactions observed between each cell type pair in CSIN. We performed these steps using all combinations of positive and negative signature genes across all cell type specific ICs (i.e. positive genes from IC1 and negative genes from IC2 and so on).

## Acknowledgements

This work is supported by the Intramural Research Program of the National Cancer Institute, Center for Cancer Research, NIH, and utilized the computational resources of the NIH HPC Biowulf cluster.

## Contributions

A.S., B.M., and V.D. curated the datasets. A.S. and B.M. developed the software for data analysis. A.S. performed most of the data analysis and developed the methodology with input from B.M and S.H. A.S. and S.H. wrote the draft of the manuscript with input from the other authors. S.H. and K.A. conceptualized this study. S.H. supervised the study with inputs from K.A.

## Notes

### Competing Interest Statement

The authors have declared no competing interest.

### Summary of Updates

We made an error when expanding the abbreviation for one of the tool that we used. The correct abbreviation for CODEFACS has bow been updated at page no. 40 as follows: CODEFACS (COnfident DEconvolution For All Cell Subset)

## References

Abdelfattah, Nourhan, Parveen Kumar, Caiyi Wang, Jia-Shiun Leu, William F. Flynn, Ruli Gao, David S. Baskin, et al. 2022. “Single-Cell Analysis of Human Glioma and Immune Cells Identifies S100A4 as an Immunotherapy Target.” Nature Communications 13(1): 767. doi:10.1038/s41467-022-28372-y.

Alshiekh Nasany, Ruham, and Macarena Ines de la Fuente. 2023. “Therapies for IDH-Mutant Gliomas.” Current Neurology and Neuroscience Reports 23(5): 225–33. doi:10.1007/s11910-023-01265-3.

Armingol, Erick, Adam Officer, Olivier Harismendy, and Nathan E. Lewis. 2021. “Deciphering Cell-Cell Interactions and Communication from Gene Expression.” Nature Reviews. Genetics 22(2): 71–88. doi:10.1038/s41576-020-00292-x.

Barkley, Dalia, Reuben Moncada, Maayan Pour, Deborah A. Liberman, Ian Dryg, Gregor Werba, Wei Wang, et al. 2022. “Cancer Cell States Recur across Tumor Types and Form Specific Interactions with the Tumor Microenvironment.” Nature Genetics 54(8): 1192–1201. doi:10.1038/s41588-022-01141-9.

Barthel, Floris P., Kevin C. Johnson, Frederick S. Varn, Anzhela D. Moskalik, Georgette Tanner, Emre Kocakavuk, Kevin J. Anderson, et al. 2019. “Longitudinal Molecular Trajectories of Diffuse Glioma in Adults.” Nature 576(7785): 112–20. doi:10.1038/s41586-019-1775-1.

Batchu, Sai, Khalid A. Hanafy, Navid Redjal, Saniya S. Godil, and Ajith J. Thomas. 2023. “Single-Cell Analysis Reveals Diversity of Tumor-Associated Macrophages and Their Interactions with T Lymphocytes in Glioblastoma.” Scientific Reports 13(1): 20874. doi:10.1038/s41598-023-48116-2.

Batlle, Eduard, and Joan Massagué. 2019. “Transforming Growth Factor-β Signaling in Immunity and Cancer.” Immunity 50(4): 924–40. doi:10.1016/j.immuni.2019.03.024.

Biton, Anne. 2024. MineICA: Analysis of an ICA Decomposition Obtained on Genomics Data.

Bremnes, Roy M., Tom Dønnem, Samer Al-Saad, Khalid Al-Shibli, Sigve Andersen, Rafael Sirera, Carlos Camps, Inigo Marinez, and Lill-Tove Busund. 2011. “The Role of Tumor Stroma in Cancer Progression and Prognosis: Emphasis on Carcinoma-Associated Fibroblasts and Non-Small Cell Lung Cancer.” Journal of Thoracic Oncology 6(1): 209–17. doi:10.1097/JTO.0b013e3181f8a1bd.

Cai, Xiangming, Feng Yuan, Junhao Zhu, Jin Yang, Chao Tang, Zixiang Cong, and Chiyuan Ma. 2021. “Glioma-Associated Stromal Cells Stimulate Glioma Malignancy by Regulating the Tumor Immune Microenvironment.” Frontiers in Oncology 11: 672928. doi:10.3389/fonc.2021.672928.

Cardoso-Moreira, Margarida, Jean Halbert, Delphine Valloton, Britta Velten, Chunyan Chen, Yi Shao, Angélica Liechti, et al. 2019. “Gene Expression across Mammalian Organ Development.” Nature 571(7766): 505–9. doi:10.1038/s41586-019-1338-5.

Chen, Binbin, Michael S. Khodadoust, Chih Long Liu, Aaron M. Newman, and Ash A. Alizadeh. 2018. “Profiling Tumor Infiltrating Immune Cells with CIBERSORT.” *Methods in Molecular Biology (Clifton*, N.J.) 1711: 243–59. doi:10.1007/978-1-4939-7493-1_12.

Cheng, Changde, Wenan Chen, Hongjian Jin, and Xiang Chen. 2023. “A Review of Single-Cell RNA-Seq Annotation, Integration, and Cell-Cell Communication.” Cells 12(15): 1970. doi:10.3390/cells12151970.

Chu, Yanshuo, Enyu Dai, Yating Li, Guangchun Han, Guangsheng Pei, Davis R. Ingram, Krupa Thakkar, et al. 2023. “Pan-Cancer T Cell Atlas Links a Cellular Stress Response State to Immunotherapy Resistance.” Nature Medicine 29(6): 1550–62. doi:10.1038/s41591-023-02371-y.

Cloughesy, Timothy F., Aaron Y. Mochizuki, Joey R. Orpilla, Willy Hugo, Alexander H. Lee, Tom B. Davidson, Anthony C. Wang, et al. 2019. “Neoadjuvant Anti-PD-1 Immunotherapy Promotes a Survival Benefit with Intratumoral and Systemic Immune Responses in Recurrent Glioblastoma.” Nature Medicine 25(3): 477–86. doi:10.1038/s41591-018-0337-7.

Colaprico, Antonio, Tiago C. Silva, Catharina Olsen, Luciano Garofano, Claudia Cava, Davide Garolini, Thais S. Sabedot, et al. 2016. “TCGAbiolinks: An R/Bioconductor Package for Integrative Analysis of TCGA Data.” Nucleic Acids Research 44(8): e71. doi:10.1093/nar/gkv1507.

Eckerdt, Frank, and Leonidas C. Platanias. 2023. “Emerging Role of Glioma Stem Cells in Mechanisms of Therapy Resistance.” Cancers 15(13): 3458. doi:10.3390/cancers15133458.

Gallus, Marco, Darwin Kwok, Senthilnath Lakshmanachetty, Akane Yamamichi, and Hideho Okada. 2023. “Immunotherapy Approaches in Isocitrate-Dehydrogenase-Mutant Low-Grade Glioma.” Cancers 15(14): 3726. doi:10.3390/cancers15143726.

Ganjoo, Shonik, Priti Gupta, Halil Ibrahim Corbali, Selene Nanez, Thomas S. Riad, Lisa K. Duong, Hampartsoum B. Barsoumian, et al. 2023. “The Role of Tumor Metabolism in Modulating T-Cell Activity and in Optimizing Immunotherapy.” Frontiers in Immunology 14: 1172931. doi:10.3389/fimmu.2023.1172931.

Gao, Ruli, Shanshan Bai, Ying C. Henderson, Yiyun Lin, Aislyn Schalck, Yun Yan, Tapsi Kumar, et al. 2021. “Delineating Copy Number and Clonal Substructure in Human Tumors from Single-Cell Transcriptomes.” Nature biotechnology 39(5): 599–608. doi:10.1038/s41587-020-00795-2.

Goldman, Mary J., Brian Craft, Mim Hastie, Kristupas Repečka, Fran McDade, Akhil Kamath, Ayan Banerjee, et al. 2020. “Visualizing and Interpreting Cancer Genomics Data via the Xena Platform.” Nature biotechnology 38(6): 675–78. doi:10.1038/s41587-020-0546-8.

Haddock, Sara, Tyler J. Alban, Şevin Turcan, Hana Husic, Eric Rosiek, Xiaoxiao Ma, Yuxiang Wang, et al. 2022. “Phenotypic and Molecular States of IDH1 Mutation-Induced CD24-Positive Glioma Stem-like Cells.” Neoplasia (New York, N.Y.) 28: 100790. doi:10.1016/j.neo.2022.100790.

Han, Yanyan, Dandan Liu, and Lianhong Li. 2020. “PD-1/PD-L1 Pathway: Current Researches in Cancer.” American Journal of Cancer Research 10(3): 727–42.

Hapke, Robert Y., and Scott M. Haake. 2020. “Hypoxia-Induced Epithelial to Mesenchymal Transition in Cancer.” Cancer Letters 487: 10–20. doi:10.1016/j.canlet.2020.05.012.

Helleday, Thomas. 2011. “The Underlying Mechanism for the PARP and BRCA Synthetic Lethality: Clearing up the Misunderstandings.” Molecular Oncology 5(4): 387–93. doi:10.1016/j.molonc.2011.07.001.

Im, Yebin, and Yongsoo Kim. 2023. “A Comprehensive Overview of RNA Deconvolution Methods and Their Application.” Molecules and Cells 46(2): 99–105. doi:10.14348/molcells.2023.2178.

Inoue, Akihiro, Takanori Ohnishi, Masahiro Nishikawa, Yoshihiro Ohtsuka, Kosuke Kusakabe, Hajime Yano, Junya Tanaka, and Takeharu Kunieda. 2023. “A Narrative Review on CD44’s Role in Glioblastoma Invasion, Proliferation, and Tumor Recurrence.” Cancers 15(19): 4898. doi:10.3390/cancers15194898.

Jerby-Arnon, Livnat, Nadja Pfetzer, Yedael Y. Waldman, Lynn McGarry, Daniel James, Emma Shanks, Brinton Seashore-Ludlow, et al. 2014. “Predicting Cancer-Specific Vulnerability via Data-Driven Detection of Synthetic Lethality.” Cell 158(5): 1199–1209. doi:10.1016/j.cell.2014.07.027.

Jerby-Arnon, Livnat, and Aviv Regev. 2022. “DIALOGUE Maps Multicellular Programs in Tissue from Single-Cell or Spatial Transcriptomics Data.” Nature Biotechnology 40(10): 1467–77. doi:10.1038/s41587-022-01288-0.

Jin, Suoqin, Christian F. Guerrero-Juarez, Lihua Zhang, Ivan Chang, Raul Ramos, Chen-Hsiang Kuan, Peggy Myung, Maksim V. Plikus, and Qing Nie. 2021. “Inference and Analysis of Cell-Cell Communication Using CellChat.” Nature Communications 12: 1088. doi:10.1038/s41467-021-21246-9.

Kelliher, Michelle A., and Justine E. Roderick. 2018. “NOTCH Signaling in T-Cell-Mediated Anti-Tumor Immunity and T-Cell-Based Immunotherapies.” Frontiers in Immunology 9: 1718. doi:10.3389/fimmu.2018.01718.

Kim, Sang Cheol, Donghyeon Yu, and Seong Beom Cho. 2018. “COEX-Seq: Convert a Variety of Measurements of Gene Expression in RNA-Seq.” Genomics & Informatics 16(4): e36. doi:10.5808/GI.2018.16.4.e36.

Korsunsky, Ilya, Nghia Millard, Jean Fan, Kamil Slowikowski, Fan Zhang, Kevin Wei, Yuriy Baglaenko, et al. 2019. “Fast, Sensitive, and Accurate Integration of Single Cell Data with Harmony.” Nature methods 16(12): 1289–96. doi:10.1038/s41592-019-0619-0.

Kotliar, Dylan, Adrian Veres, M Aurel Nagy, Shervin Tabrizi, Eran Hodis, Douglas A Melton, and Pardis C Sabeti. 2019. “Identifying Gene Expression Programs of Cell-Type Identity and Cellular Activity with Single-Cell RNA-Seq” eds. Alfonso Valencia, Naama Barkai, Elisabetta Mereu, and Berthold Göttgens. eLife 8: e43803. doi:10.7554/eLife.43803.

Kunes, Russell Z., Thomas Walle, Max Land, Tal Nawy, and Dana Pe’er. 2024. “Supervised Discovery of Interpretable Gene Programs from Single-Cell Data.” Nature Biotechnology 42(7): 1084–95. doi:10.1038/s41587-023-01940-3.

Lee, Joo Sang, Avinash Das, Livnat Jerby-Arnon, Rand Arafeh, Noam Auslander, Matthew Davidson, Lynn McGarry, et al. 2018. “Harnessing Synthetic Lethality to Predict the Response to Cancer Treatment.” Nature Communications 9: 2546. doi:10.1038/s41467-018-04647-1.

von Locquenghien, Michelle, Catalina Rozalén, and Toni Celià-Terrassa. “Interferons in Cancer Immunoediting: Sculpting Metastasis and Immunotherapy Response.” The Journal of Clinical Investigation 131(1): e143296. doi:10.1172/JCI143296.

Luca, Bogdan A., Chloé B. Steen, Magdalena Matusiak, Armon Azizi, Sushama Varma, Chunfang Zhu, Joanna Przybyl, et al. 2021. “Atlas of Clinically Distinct Cell States and Ecosystems across Human Solid Tumors.” Cell 184(21): 5482–5496.e28. doi:10.1016/j.cell.2021.09.014.

Madrigal, Pedro, Siwei Deng, Yuliang Feng, Stefania Militi, Kim Jee Goh, Reshma Nibhani, Rodrigo Grandy, et al. 2023. “Epigenetic and Transcriptional Regulations Prime Cell Fate before Division during Human Pluripotent Stem Cell Differentiation.” Nature Communications 14(1): 405. doi:10.1038/s41467-023-36116-9.

Magen, Assaf, Avinash Das Sahu, Joo Sang Lee, Mahfuza Sharmin, Alexander Lugo, J. Silvio Gutkind, Alejandro A. Schäffer, Eytan Ruppin, and Sridhar Hannenhalli. 2019. “Beyond Synthetic Lethality: Charting the Landscape of Pairwise Gene Expression States Associated with Survival in Cancer.” Cell reports 28(4): 938–948.e6. doi:10.1016/j.celrep.2019.06.067.

Mansouri, Siavash, Daniel Heylmann, Thorsten Stiewe, Michael Kracht, and Rajkumar Savai. 2022. “Cancer Genome and Tumor Microenvironment: Reciprocal Crosstalk Shapes Lung Cancer Plasticity.” eLife 11: e79895. doi:10.7554/eLife.79895.

Martyshkina, Yuliya S., Valeriy P. Tereshchenko, Daria A. Bogdanova, and Stanislav A. Rybtsov. 2023. “Reliable Hallmarks and Biomarkers of Senescent Lymphocytes.” International Journal of Molecular Sciences 24(21): 15653. doi:10.3390/ijms242115653.

Matlak, Dariusz, and Ewa Szczurek. 2017. “Epistasis in Genomic and Survival Data of Cancer Patients.” PLoS computational biology 13(7): e1005626. doi:10.1371/journal.pcbi.1005626.

Miller, Julie J., Franziska Loebel, Tareq A. Juratli, Shilpa S. Tummala, Erik A. Williams, Tracy T. Batchelor, Isabel Arrillaga-Romany, and Daniel P. Cahill. 2019. “Accelerated Progression of IDH Mutant Glioma after First Recurrence.” Neuro-Oncology 21(5): 669–77. doi:10.1093/neuonc/noz016.

Müller, Sören, Ara Cho, Siyuan J Liu, Daniel A Lim, and Aaron Diaz. 2018. “CONICS Integrates scRNA-Seq with DNA Sequencing to Map Gene Expression to Tumor Sub-Clones.” Bioinformatics 34(18): 3217–19. doi:10.1093/bioinformatics/bty316.

Murata, Akihiko, Miya Yoshino, Mari Hikosaka, Kazuki Okuyama, Lan Zhou, Seiji Sakano, Hideo Yagita, and Shin-Ichi Hayashi. 2014. “An Evolutionary-Conserved Function of Mammalian Notch Family Members as Cell Adhesion Molecules.” PloS One 9(9): e108535. doi:10.1371/journal.pone.0108535.

Nawy, Tal. 2018. “A Pan-Cancer Atlas.” Nature Methods 15(6): 407–407. doi:10.1038/s41592-018-0020-4.

Neftel, Cyril, Julie Laffy, Mariella G. Filbin, Toshiro Hara, Marni E. Shore, Gilbert J. Rahme, Alyssa R. Richman, et al. 2019. “An Integrative Model of Cellular States, Plasticity, and Genetics for Glioblastoma.” Cell 178(4): 835–849.e21. doi:10.1016/j.cell.2019.06.024.

O’Neil, Nigel J., Melanie L. Bailey, and Philip Hieter. 2017. “Synthetic Lethality and Cancer.” Nature Reviews Genetics 18(10): 613–23. doi:10.1038/nrg.2017.47.

Phillips, Patrick C. 2008. “Epistasis--the Essential Role of Gene Interactions in the Structure and Evolution of Genetic Systems.” Nature Reviews. Genetics 9(11): 855–67. doi:10.1038/nrg2452.

Qazi, M. A., P. Vora, C. Venugopal, S. S. Sidhu, J. Moffat, C. Swanton, and S. K. Singh. 2017. “Intratumoral Heterogeneity: Pathways to Treatment Resistance and Relapse in Human Glioblastoma.” Annals of Oncology: Official Journal of the European Society for Medical Oncology 28(7): 1448–56. doi:10.1093/annonc/mdx169.

Qu, Yidi, Bo Dou, Horyue Tan, Yibin Feng, Ning Wang, and Di Wang. 2019. “Tumor Microenvironment-Driven Non-Cell-Autonomous Resistance to Antineoplastic Treatment.” Molecular Cancer 18(1): 69. doi:10.1186/s12943-019-0992-4.

Ramirez, Christel F. A., Daniel Taranto, Masami Ando-Kuri, Marnix H. P. de Groot, Efi Tsouri, Zhijie Huang, Daniel de Groot, et al. 2024. “Cancer Cell Genetics Shaping of the Tumor Microenvironment Reveals Myeloid Cell-Centric Exploitable Vulnerabilities in Hepatocellular Carcinoma.” Nature Communications 15: 2581. doi:10.1038/s41467-024-46835-2.

Ravi, Vidhya M., Nicolas Neidert, Paulina Will, Kevin Joseph, Julian P. Maier, Jan Kückelhaus, Lea Vollmer, et al. 2022. “T-Cell Dysfunction in the Glioblastoma Microenvironment Is Mediated by Myeloid Cells Releasing Interleukin-10.” Nature Communications 13(1): 925. doi:10.1038/s41467-022-28523-1.

Reis, Bernardo S., Patrick W. Darcy, Iasha Z. Khan, Christine S. Moon, Adam E. Kornberg, Vanessa S. Schneider, Yelina Alvarez, et al. 2022. “TCR-Vγδ Usage Distinguishes Protumor from Antitumor Intestinal Γδ T Cell Subsets.” *Science (New York*, N.Y.) 377(6603): 276–84. doi:10.1126/science.abj8695.

Reuss, David. E. 2023. “Updates on the WHO Diagnosis of IDH-Mutant Glioma.” Journal of Neuro-Oncology 162(3): 461–69. doi:10.1007/s11060-023-04250-5.

Sharma, Pratibha, Ashley Aaroe, Jiyong Liang, and Vinay K. Puduvalli. 2023. “Tumor Microenvironment in Glioblastoma: Current and Emerging Concepts.” Neuro-Oncology Advances 5(1): vdad009. doi:10.1093/noajnl/vdad009.

Singh, Arashdeep, Arati Rajeevan, Vishaka Gopalan, Piyush Agrawal, Chi-Ping Day, and Sridhar Hannenhalli. 2022. “Broad Misappropriation of Developmental Splicing Profile by Cancer in Multiple Organs.” Nature Communications 13(1): 7664. doi:10.1038/s41467-022-35322-1.

Sompairac, Nicolas, Petr V. Nazarov, Urszula Czerwinska, Laura Cantini, Anne Biton, Askhat Molkenov, Zhaxybay Zhumadilov, et al. 2019. “Independent Component Analysis for Unraveling the Complexity of Cancer Omics Datasets.” International Journal of Molecular Sciences 20(18): 4414. doi:10.3390/ijms20184414.

Sorrelle, Noah, Adrian T. A. Dominguez, and Rolf A. Brekken. 2017. “From Top to Bottom: Midkine and Pleiotrophin as Emerging Players in Immune Regulation.” Journal of Leukocyte Biology 102(2): 277–86. doi:10.1189/jlb.3MR1116-475R.

Stuart, Tim, Andrew Butler, Paul Hoffman, Christoph Hafemeister, Efthymia Papalexi, William M. Mauck, Yuhan Hao, et al. 2019. “Comprehensive Integration of Single-Cell Data.” Cell 177(7): 1888–1902.e21. doi:10.1016/j.cell.2019.05.031.

Szczurek, Ewa, Navodit Misra, and Martin Vingron. 2013. “Synthetic Sickness or Lethality Points at Candidate Combination Therapy Targets in Glioblastoma.” International Journal of Cancer 133(9): 2123–32. doi:10.1002/ijc.28235.

Tirosh, Itay, Andrew S. Venteicher, Christine Hebert, Leah E. Escalante, Anoop P. Patel, Keren Yizhak, Jonathan M. Fisher, et al. 2016. “Single-Cell RNA-Seq Supports a Developmental Hierarchy in Human Oligodendroglioma.” Nature 539(7628): 309–13. doi:10.1038/nature20123.

del Toro, Raquel, Claudia Prahst, Thomas Mathivet, Geraldine Siegfried, Joshua S. Kaminker, Bruno Larrivee, Christiane Breant, et al. 2010. “Identification and Functional Analysis of Endothelial Tip Cell-Enriched Genes.” Blood 116(19): 4025–33. doi:10.1182/blood-2010-02-270819.

van der Vaart, Thijs, Maarten M.J. Wijnenga, Karin van Garderen, Hendrikus J. Dubbink, Pim J. French, Marion Smits, Clemens M.F. Dirven, et al. 2024. “Differences in the Prognostic Role of Age, Extent of Resection, and Tumor Grade between Astrocytoma IDHmt and Oligodendroglioma: A Single-Center Cohort Study.” Clinical Cancer Research 30(17): 3837–44. doi:10.1158/1078-0432.CCR-24-0901.

Venteicher, Andrew S., Itay Tirosh, Christine Hebert, Keren Yizhak, Cyril Neftel, Mariella G. Filbin, Volker Hovestadt, et al. 2017. “Decoupling Genetics, Lineages, and Microenvironment in IDH-Mutant Gliomas by Single-Cell RNA-Seq.” *Science (New York*, N.Y.) 355(6332): eaai8478. doi:10.1126/science.aai8478.

de Visser, Karin E., and Johanna A. Joyce. 2023. “The Evolving Tumor Microenvironment: From Cancer Initiation to Metastatic Outgrowth.” Cancer Cell 41(3): 374–403. doi:10.1016/j.ccell.2023.02.016.

de Vries, Natasja L., Ahmed Mahfouz, Frits Koning, and Noel F. C. C. de Miranda. 2020. “Unraveling the Complexity of the Cancer Microenvironment With Multidimensional Genomic and Cytometric Technologies.” Frontiers in Oncology 10: 1254. doi:10.3389/fonc.2020.01254.

Wang, Kun, Sushant Patkar, Joo Sang Lee, E. Michael Gertz, Welles Robinson, Fiorella Schischlik, David R. Crawford, Alejandro A. Schäffer, and Eytan Ruppin. 2022. “Deconvolving Clinically Relevant Cellular Immune Crosstalk from Bulk Gene Expression Using CODEFACS and LIRICS Stratifies Melanoma Patients to Anti-PD-1 Therapy.” Cancer discovery 12(4): 1088–1105. doi:10.1158/2159-8290.CD-21-0887.

Wechter, Noah, Martina Rossi, Carlos Anerillas, Dimitrios Tsitsipatis, Yulan Piao, Jinshui Fan, Jennifer L. Martindale, et al. 2023. “Single-Cell Transcriptomic Analysis Uncovers Diverse and Dynamic Senescent Cell Populations.” Aging 15(8): 2824–51. doi:10.18632/aging.204666.

Whitfield, Benjamin T., and Jason T. Huse. 2022. “Classification of Adult-Type Diffuse Gliomas: Impact of the World Health Organization 2021 Update.” Brain Pathology (Zurich, Switzerland) 32(4): e13062. doi:10.1111/bpa.13062.

Woroniecka, Karolina I., Kristen E. Rhodin, Pakawat Chongsathidkiet, Kristin A. Keith, and Peter E. Fecci. 2018. “T-Cell Dysfunction in Glioblastoma: Applying a New Framework.” Clinical Cancer Research: An Official Journal of the American Association for Cancer Research 24(16): 3792–3802. doi:10.1158/1078-0432.CCR-18-0047.

Xia, Longzheng, Linda Oyang, Jinguan Lin, Shiming Tan, Yaqian Han, Nayiyuan Wu, Pin Yi, et al. 2021. “The Cancer Metabolic Reprogramming and Immune Response.” Molecular Cancer 20(1): 28. doi:10.1186/s12943-021-01316-8.

Zhang, Jiayu, Tong Lu, Shiqi Lu, Shuaijun Ma, Donghui Han, Keying Zhang, Chao Xu, et al. 2022. “Single-Cell Analysis of Multiple Cancer Types Reveals Differences in Endothelial Cells between Tumors and Normal Tissues.” Computational and Structural Biotechnology Journal 21: 665–76. doi:10.1016/j.csbj.2022.12.049.

Zhang, Yang, Yang Liu, Fengchao Lang, and Chunzhang Yang. 2022. “IDH Mutation and Cancer Stem Cell.” Essays in Biochemistry 66(4): 413–22. doi:10.1042/EBC20220008.

Zhao, Zheng, Ke-Nan Zhang, Qiangwei Wang, Guanzhang Li, Fan Zeng, Ying Zhang, Fan Wu, et al. 2021. “Chinese Glioma Genome Atlas (CGGA): A Comprehensive Resource with Functional Genomic Data from Chinese Glioma Patients.” Genomics, Proteomics & Bioinformatics 19(1): 1–12. doi:10.1016/j.gpb.2020.10.005.

